# Insight into GABA shunt-associated aldehyde dehydrogenases (ALDH) and stress responses of ALDH superfamily in moss and barley

**DOI:** 10.64898/2026.01.13.699213

**Authors:** David J. Kopečný, Jakub Bělíček, Martina Kopečná, Armelle Vigouroux, Radka Končitíková, Klaus von Schwartzenberg, Klára Končáková, Sanja Ćavar Zeljković, Miroslav Valárik, Karel Müller, Roman Kouřil, Veronique Bergougnoux, Solange Moréra, David Kopečný

**Affiliations:** Department of Experimental Biology, Faculty of Science, Palacký University, Olomouc CZ-77900, Czech Republic; Université Paris-Saclay, CEA, CNRS, Institute for Integrative Biology of the Cell (I2BC), Gif-sur-Yvette F-91198, France; Institute for Plant Science and Microbiology, Universität Hamburg, D-22609 Hamburg, Germany; Czech Advanced Technology and Research Institute, Palacký University, CZ-77900 Olomouc, Czech Republic; Czech Agrifood Research Center, Olomouc 77900, Czech Republic; Centre of Plant Structural and Functional Genomics, Institute of Experimental Botany of the Czech Academy of Sciences, CZ-77900 Olomouc, Czech Republic; Transcriptomics and Cloning Facility, Institute of Experimental Botany of the Czech Academy of Sciences, CZ-16502 Prague, Czech Republic; Department of Biophysics, Faculty of Science, Palacký University, Olomouc CZ-77900, Czech Republic

**Keywords:** Abiotic stress, aldehyde dehydrogenase, GABA shunt, glutamate, glutathione-S-transferase, *Hordeum vulgare*, hormone, knockout, *Physcomitrium patens*, succinic semialdehyde, X-ray structure

## Abstract

We explored the expression of the aldehyde dehydrogenase (ALDH) superfamily in two model plants, *Physcomitrium patens* (moss) and *Hordeum vulgare* (barley), under various stress conditions. The ALDH enzymes are crucial for oxidizing aldehydes to carboxylic acids and are involved in multiple metabolic pathways. We found significant differences in enzyme expression between moss and barley within the same ALDH families. We then focused on the ALDH5, ALDH10, and ALDH21 families, which are part of the γ-aminobutyric acid (GABA) shunt, noting that the ALDH21 family is absent in barley. The kinetic properties of ALDH10 and ALDH5 enzymes were analyzed, revealing that PpALDH5F1 exhibits high specificity for succinic semialdehyde (SSAL), a product of GABA. The crystal structure of PpALDH5F1 identified key residues for SSAL binding. Knockout mutants of moss *aldh5F2, aldh10A1, and aldh21A1* showed slightly smaller colonies than the wild-type. GABA and glutamate levels were elevated in *aldh5F2* and *aldh21A1* knockouts due to a partially blocked GABA shunt pathway, while *aldh10A1* knockout showed no changes in GABA levels. Transcriptomic data revealed a link between several genes, including six upregulated *glutathione-S-transferase* genes in all three *aldh* knockouts, suggesting a direct compensatory mechanism for oxidative stress protection via conjugation of undegraded aldehydes to glutathione.

**Highlight:** Moss knockouts of GABA shunt-associated aldehyde dehydrogenases display slower growth, changes in levels of glutamate, glutamine and GABA, and result in upregulation of several unique *glutathione-S-transferase* genes.

## INTRODUCTION

ALDHs constitute a superfamily of NAD(P)^+^-dependent enzymes that detoxify released aldehydes to nontoxic carboxylic acids and at least 13 ALDH families have been identified in plants (Brocker *et al*., 2013; Tola *et al*., 2020; Stiti *et al*., 2021). ALDHs sharing more than 40% sequence identity belong to the same family. Each family number is followed by a letter representing the given subfamily and another number for the individual gene within that subfamily. Many of the plant *ALDH* genes are stress-responsive, indicating their importance in adaptation to environmental changes (Sunkar *et al*., 2003). ALDHs are associated with various metabolic pathways, as well as metabolites that play a crucial role in stress responses. For example, reactive oxygen species are responsible for the peroxidation of membrane lipids and the production of short aliphatic aldehydes such as malondialdehyde, hexanal, nonanal and their unsaturated and hydroxylated derivatives, which can be detoxified by members of ALDH2 and ALDH3 families (Končitíková *et al*., 2015; Sunkar *et al*., 2003; Stiti *et al*., 2011). Additionally, the coenzyme product NAD(P)H contributes to the maintenance of cellular redox balance.

Land plants sense the nutrient and energy state of the cells and perform metabolic and developmental adaptations using, for example, “Target of Rapamycin” and “AMP-regulated kinase/Sucrose non-fermenting 1” kinase pathways (Robaglia *et al.,* 2012). The contribution of hormones is also important. Abscisic acid (ABA) plays a crucial role in osmotic stress responses, while ethylene is involved in submergence responses (Toriyama *et al*., 2022; Yasumura *et al*., 2012). Among other hormones, sensing of cytokinin and auxin, key factors in plant growth, is already present in mosses (von Schwartzenberg K *et al*., 2016; Thelander *et al*., 2018). Jasmonate signaling is slightly different in mosses, as only the cyclopentenone precursor of jasmonic acid is synthesized (Stumpe *et al*., 2010; Monte *et al*., 2020). Nevertheless, mosses respond to methyl jasmonate (Ponce de León *et al*., 2012).

Stress adaptation includes the production of compatible osmolytes or changes in nitrogen-rich compounds. Proline acts both as a compatible osmolyte, which accumulates during abiotic stresses, and as an energy source during pollen maturation and germination in plants (Hare and Cress, 1997). Plant ALDH12 regulates the degradation of proline (Pro) as well as of ornithine and arginine (Arg) into glutamate (Glu) (Korasick *et al*., 2019), while the bi-functional ALDH18 catalyzes the synthesis of Pro from Glu (Funck *et al*., 2020). Arg is a storage form of organic nitrogen and a precursor for polyamine synthesis (Bagni and Tassoni, 2001). Polyamines are important secondary metabolites and for example, spermidine is well-known to activate the plant antioxidative responses under abiotic stress (Seifi and Shelp, 2019).

Catabolism of polyamines, Arg or lysine releases highly reactive aldehydes such as 3-aminopropionaldehyde (APAL), 4-aminobutyraldehyde (ABAL), trimethyl-4-aminobutyraldehyde (TMABAL) and 4-guanidinobutyraldehyde (GBAL), which are all detoxified by members of the ALDH10 family (also annotated as aminoaldehyde dehydrogenase, AMADH) (Tylichová *et al*., 2010; Kopečný *et al*., 2013). Plant ALDH7 contributes to lysine catabolism (Končitíková *et al*., 2015). Several ALDH10 reaction products, such as glycine betaine (GB), β-alanine or trimethyl-4-aminobutyrate (TMABA), function as compatible osmolytes (Le Rudulier *et al*., 1984; Rhodes and Hanson, 1993; Duhazé *et al*., 2003; Jacques *et al*., 2020). Oxidation of ABAL leads to γ-aminobutyric acid (GABA), which is involved in numerous processes (Michaeli and Fromm, 2015; Shelp *et al*., 2021). GABA reduces stomatal opening and consequently loss of water (Xu *et al*., 2021), affects pollen development (Palanivelu *et al*., 2003), modulates the activity of 14-3-3 proteins and carbon-nitrogen balance (Lancien and Roberts, 2006; Fait *et al*., 2011), and participates in biotic responses via the 2-hexenal signaling pathway (Mirabella *et al*., 2008). GABA also acts as a signaling molecule by regulating aluminum-activated malate transporter channels, which play a key role in fine-tuning responses to abiotic stress (Ramesh *et al*., 2015).

Notably, GABA interconnects the polyamine degradation with the GABA shunt, which is considered an adaptive metabolic pathway that helps to maintain the respiratory rate in mitochondria by supplying succinate to the tricarboxylic acid (TCA) cycle (Studart-Guimarães *et al*., 2007). The GABA shunt begins in the cytosol, where GABA is synthesized either from Glu by the Glu decarboxylase (GAD) or from polyamines by the poly- and diamine oxidases (PAOs and DAOs) and ALDH10 (Fig. 1). GABA is transported into the mitochondria by GABA permease (Michaeli *et al*., 2011), where it is converted to succinic semialdehyde (SSAL) by GABA transaminase (GABA-T). Subsequently, SSAL is either oxidized to succinate by SSAL dehydrogenases (often abbreviated as SSADHs or SSALDHs) or reduced to 4-hydroxybutyrate (GHB) by a glyoxylate reductase (Michaeli and Fromm, 2015; Shelp *et al*., 2021). The succinate is further oxidized by succinate dehydrogenase, which also serves as an entry point to the electron transport chain. SSALDHs comprise two distinct ALDH families: the mitochondrial ALDH5 uses NAD^+^ as a coenzyme and the cytosolic ALDH21 utilizes NADP^+^. The latter family is absent in higher plants and has been characterized in detail in *Physcomitrium* (Kopečná *et al*., 2017).

**Fig. 1.**
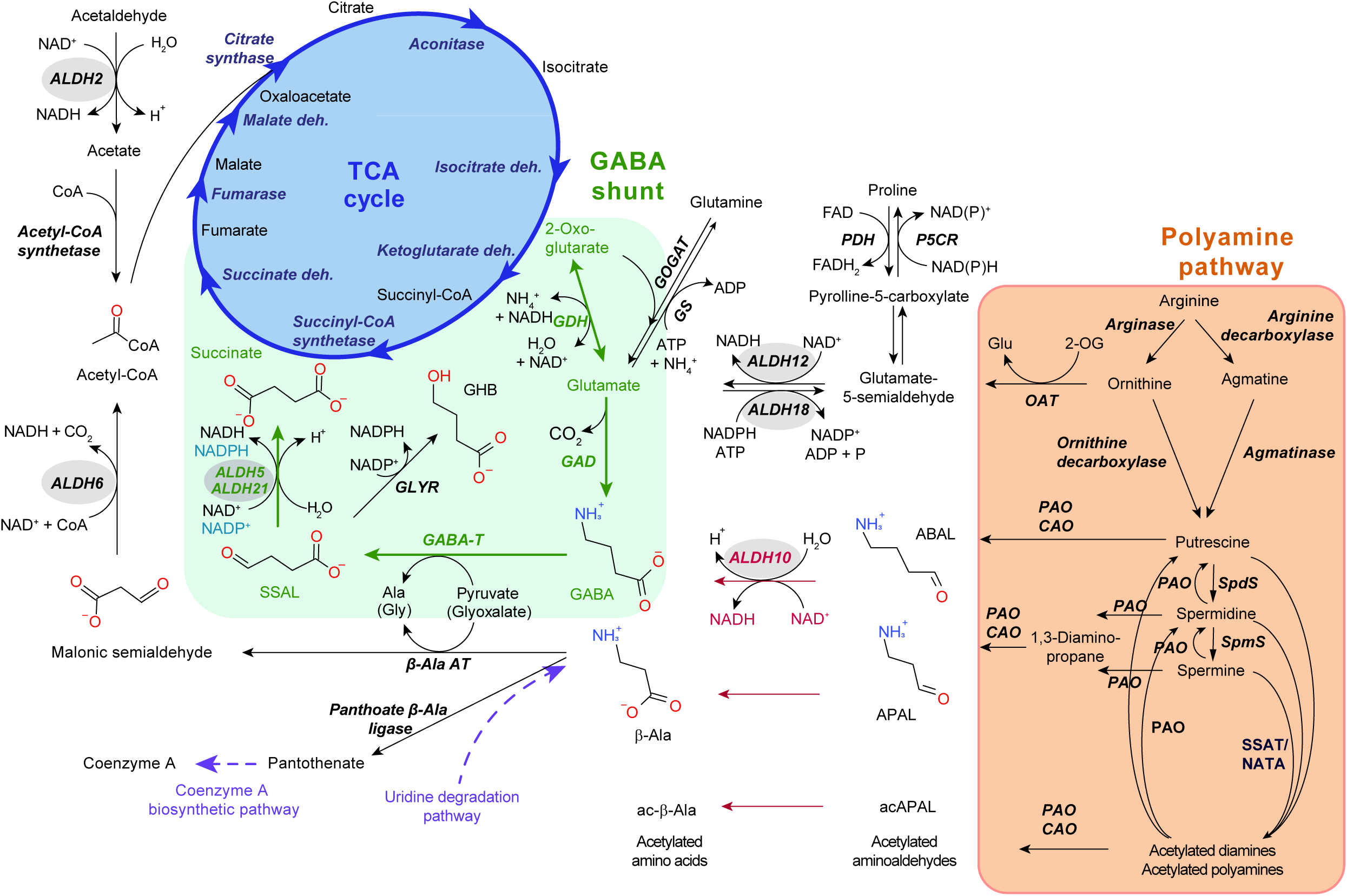
Reaction scheme of the TCA cycle and its connection to the GABA shunt and polyamine pathway via ALDH10 and ALDH5 reactions. GABA, which is synthesized from glutamate or 4-aminobutyrate (ABAL), is transaminated via GABA-T to succinic semialdehyde (SSAL) and further oxidized to succinate by plant ALDH5 and ALDH21 using NAD^+^ and NADP^+^ as coenzymes, respectively. NAD^+^-dependent ALDH10 interconnects GABA shunt (green) and polyamine metaboli-sm (orange) as it oxidizes 3-aminopropanal (APAL), 3-acetamidopropanal (acAPAL) and ABAL from di-/poly-amine degra-dation to amino acids such as *β*-Ala, ac-*β*-Ala and GABA.

Here, we analyzed the expression pattern of the *ALDH* superfamily in two model plants, namely *Physcomitrium patens* and *Hordeum vulgare*. *Physcomitrium* is a highly desiccation-tolerant nonvascular moss that diverged from unicellular algae more than 400 million years ago and is a close living relative of the earliest land plants (Rensing *et al*., 2008). *Hordeum vulgare* cultivar Morex is a US spring six-row malting barley (Mascher *et al*., 2021), one of the most economically important crops worldwide, representing a model of the higher plant. We analyzed how abiotic stress responses were altered among the same *ALDH* families in these two model plants. In particular, we focused on families related to the GABA shunt pathway. While the moss ALDH5 and ALDH10 enzymes have not been studied yet, those from barley have been reported (Yamaura *et al*., 1988; Fujiwara *et al*., 2008). We analyzed the kinetic properties of the single PpALDH10 and two PpALDH5 members. We solved the first plant crystal structure of PpALDH5F1, revealing the binding mode of SSAL. Because mosses possess a unique drought-inducible *ALDH21*, absent in barley, we investigated whether *ALDH21* confers specific benefits under stress and whether related ALDHs can compensate each other. Therefore, we produced moss *aldh5F2*, *aldh10A1* and *aldh21A1* knockouts to obtain complementary information on the importance of each family for the regulation of GABA levels and analyzed levels of closely related metabolites.

## MATERIALS AND METHODS

### Barley (Hordeum vulgare) and moss (Physcomitrium patens) genome analyses

A genome of barley *Hordeum vulgare* L. cv. Morex (MorexV3) and moss *Physcomitrium patens* v6.1 (Mascher *et al*., 2021; Bi *et al*., 2024) at Ensembl Plants (https://plants.ensembl.org/index.html) and Phytozome (https://phytozome-next.jgi.doe.gov/) were screened for *ALDH* genes. Regions of the genome with significant identity, including 2000 bp before the start and after the stop codon, were retrieved using the Samtools-faidx function from the Samtools package (Li *et al*., 2009). The regions with 100% identity were gapped due to the possible presence of introns. The gene regions were manually reannotated and compared to the RNA sequences deposited at NCBI using the blastn tool (https://blast.ncbi.nlm.nih.gov/Blast.cgi). RNAs with 98-100 % identity and E value below 1e^-5^ were considered.

### Quantitative PCR (qPCR) analysis of ALDH expression in barley and moss

Moss *P. patens* (‘Gransden 2004‘ strain) was grown in liquid Knop’s medium (15 mM nitrogen) and then exposed for 48 hours either to 400 mM mannitol, 300 mM NaCl, or nitrogen stress (N-stress, 1.5 mM nitrogen; N+ stress, 24 mM nitrogen as calcium nitrate). Hormone treatment with 10 µM abscisic acid (ABA), methyl jasmonate (MeJA), cytokinin derivative benzyl adenine (BA) or auxin analog 2,4-dichlorophenoxyacetic acid (2,4-D) lasted for 24 hours. Heat stress was done at 37 °C for 1 hour, followed by a recovery period; cold stress was done for 5 hours at 5 °C. Two-week-old barley (cv. Golden Promise) seedlings were grown hydroponically in standard Hoagland solution (15 mM nitrogen; ammonium nitrate) and then exposed to osmotic stress (20 % PEG6000) or salt stress (200 mM NaCl) for 48 hours. Nitrogen and hormone treatments were identical to those of moss.

Plant material, either two 200 ml moss cultures or a pool of 20 barley seedlings, was disrupted in a mixer mill (MM 200; www.retsch.com). The RNA from moss was extracted using the citrate/citric acid method (Oñate-Sánchez and Vicente-Carbajosa, 2008). The final samples were then precipitated with LiCl, washed twice with 70% ethanol (v/v), and diluted in DEPC-treated water. The RNA was extracted from barley seedlings using the phenol-chloroform method with TRIzol reagent and treated with Turbo DNase (www.thermofisher.com). Transcription was performed in duplicate using a LunaScript RT SuperMix, and qPCR was conducted in six technical replicates using an RT Luna Universal Probe qPCR Master Mix (www.neb.com) on a QuantStudio 5 Real-Time PCR System (www.thermofisher.com). Primers and the FAM-TAM probes were designed using Primer Express 3.0 software. They are listed in Supplementary Table S1 and Table S2. Plasmid constructs carrying the *PpALDH* ORFs were used for PCR efficiency determination. Cycle threshold values were normalized to moss *elongation factor 1*α (Pp6c2_4070) and *ubiquitin 2* (Pp6c14_11780), or barley *elongation factor 2* (HORVU.MOREX.r3.5HG0529120) and β*-actin* (HORVU.MOREX.r3.2HG0099270) genes and amplification efficiency.

### Promoter DNA sequence analysis

For each gene, the 1,000 bp region upstream of the start codon was analyzed for plant cis-acting regulatory elements using the PlantCARE database (Lescot *et al*., 2002, http://bioinformatics.psb.ugent.be/webtools/plantcare/html/).

### Cloning, expression and purification of selected ALDHs

The total RNA from barley leaves and *Physcomitrium patens* at the protonema stage was extracted using the RNAqueous kit and Plant RNA Isolation Aid. The cDNA was synthesized using Superscript IV RT and oligo dT primers (https://www.thermofisher.com). The sequences coding for PpALDH5F2 (Pp6c6_14560V6.1, 1497 bp) and PpALDH5F1 (Pp6c26_6240V6.1, 1479 bp) were amplified (Supplementary Table S3) by the Q5 High-Fidelity DNA polymerase (www.neb.com). *HvALDH5F1* (HORVU6Hr1G031480.3, 1581 bp) was amplified to obtain Δ_1-32_HvALDH5F1 devoid of the putative mitochondrial signal sequence, while *PpALDH10A1* (Pp6c5_11700V6.1, 1680 bp) was amplified to obtain Δ_1-52_PpALDH10A1 devoid of the putative chloroplastic signal sequence (Supplementary Table S3). The amplicons were further cloned into a pET28b vector (www.merckmillipore.com) and then transformed into Rosetta 2 (DE3) *E. coli* cells (www.merckmillipore.com). Mutants of *PpALDH5F1* were cloned using phosphorylated tail-to-tail oriented primers (Supplementary Table S3), with the mutation being located at the 5’ end of one of the primers by a PCR reaction in 35 cycles using AccuPrime *Pfx* polymerase (https://www.thermofisher.com). The products were treated with the *Dpn*I, gel-purified and ligated using the T4 DNA ligase (https://www.thermofisher.com) and transformed into Rosetta 2 (DE3) *E. coli* cells. Protein production was carried out at 20 °C overnight using 0.5 mM IPTG. ALDHs were purified on an NGC Medium-Pressure Liquid Chromatography System (https://www.bio-rad.com) on a Nickel-HiTrap IMAC FF column and further on a HiLoad 26/60 Superdex 200 column (https://www.cytivalifesciences.com) in 20 mM Tris-HCl buffer, pH 8.0, 100 mM NaCl. We obtained 6 to 11 mg of pure proteins from 200 ml cultures. Gel permeation chromatography analyses were performed on a Superdex 200 10/30 HR column in 50 mM phosphate buffer with 100 mM NaCl using Gel Filtration Standard (Bio-Rad).

### Enzyme kinetics

The enzyme activity was measured by monitoring the NAD(P)H formation (ε_340_ = 6.22 mM^−1^ cm^−1^) at 30 °C on an Agilent UV-Vis spectrophotometer 8453 (www.agilent.com) in 150 mM HEPES buffer at pH 8.2 and 1.5 mM NAD^+^. Substrate screening was done using 1 mM aldehydes for ALDH10 and 0.25 mM aldehydes for ALDH5. Saturation curves for NAD^+^ and NADP^+^ coenzymes were measured using SSAL concentration, providing the highest rates, i.e. 200 μM for PpALDH5F2 and 40 μM for PpALDH5F1/HvALDH5F1. SSAL was purchased from Santa Cruz Biotechnology (www.scbt.com) and AASAL ethylene acetal was from Chiralix (www.chiralix.com). BAL chloride, elementary aliphatic aldehydes, or APAL and ABAL diethyl acetals were purchased from Sigma-Aldrich (www.sigmaaldrich.com) and GRSAL was synthesized by oxidation of D,L-2-aminoadipic acid with chloramine-T (Adams and Chan, 1971). APAL, ABAL, GBAL, TMABAL or AASAL were always freshly prepared (Tylichová *et al*., 2010). Kinetic constants, including their S.E. values, were determined by nonlinear regression using the GraphPad Prism 8.0 software (www.graphpad.com).

### Crystallization and structure determination

Crystallization conditions of purified PpALDH5F1 at 10.0 mg ml^-1^ were found with the Classic Suite (www.qiagen.com). Crystals of the apoform or the complex with 2 mM NAD^+^ were obtained in hanging drops in the presence of 1.7 M LiSO_4_ in 100 mM HEPES, pH 7.5. Those with NAD^+^ comprised 100 mM sodium succinate. Crystals were cryoprotected in mother liquor supplemented with 18% PEG 400 or 25% glycerol and flash-frozen in liquid nitrogen. Diffraction data were collected at 100 K on the PROXIMA 1 and 2 beamlines at the SOLEIL synchrotron (www.synchrotron-soleil.fr). Intensities were integrated using the autoPROC (www.globalphasing.com) and XDS program (Kabsch, 2010) and further reprocessed by STARANISO (Tickle *et al*., 2018). Molecular replacement was performed with Phaser (Storoni *et al*., 2004) using the structure of the *Homo sapiens* ALDH5 (PDB 2W8Q) (Kim *et al*., 2009) as the search model. Models were refined with NCS restraints and TLS using Buster 2.10 (Bricogne *et al*., 2011) and with ligand occupancies set to 1. Electron density maps were evaluated using COOT (Emsley and Cowtan, 2004). Molecular graphics images were generated using PYMOL v 2.5 (www.pymol.org). Data collection and refinement statistics are in Supplementary Table S4.

### Affinity and thermal stability determination

The MST and spectral shift methods were used to determine the binding affinity using Monolith NT.115 and Monolith X instruments (www.nanotemper-technologies.com). The enzymes were fluorescently labeled using the RED-tris-NTA dye (2:1 dye/protein molar ratio). Labeled PpALDH10 was adjusted to 100 nM with 50 mM Tris-HCl buffer, 100 mM NaCl and 1% GOL. Measurements were performed in premium capillaries at 25 °C. Thermal stability was measured using nano-differential scanning fluorimetry on Tycho NT.6 and Prometheus Panta instruments (www.nanotemper-technologies.com) in various buffers and in the presence of putative substrates and coenzyme, in a range of 20 to 95 °C, with a heating rate of 1 °C min^-1^, and using Panta control software. Protein unfolding was measured by detecting the temperature-induced change in tryptophan fluorescence intensity at emission wavelengths of 330 and 350 ± 5 nm. The melting temperature (*T*_m_) was deduced from the maximum of the first derivative of the fluorescence ratios F_350_/F_330_.

### Cloning of gene replacement vectors and preparation of Physcomitrium knockout mutants

Functional *aldh5F2*, *aldh10A1*, and *aldh21A1* moss knockouts were prepared using a gene-replacement pBNRF vector containing a resistance cassette flanked by 1000- to 1200-bp-long genomic fragments from the 5′ and 3′ regions of the corresponding genomic *PpALDH* loci. The fragments were cloned (Supplementary Table S5), further digested and ligated into the pBNRF vector, conferring G418 resistance. The transformation of *P. patens* protoplasts (Schaefer *et al*., 1991) was carried out using the constructed replacement vector, from which the respective transformation cassette was released upon double digest. Up to 100 stable moss lines were obtained after three rounds of selection. The haploid status of ∼ 50 transformants was verified by flow cytometry and up to 20 transgenic lines for each mutant were then analyzed by PCR for homologous recombination at the corresponding loci and by qPCR for the absence of the transcript using FAM-TAMRA probes. Finally, seven independent homozygous *aldh5F2* lines, eight *aldh10A1* lines and five *aldh21A1* lines were selected. Strains were grown in a BCD medium and maintained in growth chambers at 25 °C illuminated with white light under 16/8-h light/dark periods with a photon flux of 50 µmol m^−2^ s^−1^.

### Phenotyping and metabolite screening in moss upon drought and salt stresses

Three *aldh* knockout lines for each gene were chosen for metabolite and phenotypic characterization in *P. patens*. Point inoculation was performed with three independent lines for each knockout into 6- and 24-well plates containing modified Knop medium (Reski and Abel, 1985) enriched with trace element solution, vitamins and tartrate and NaCl (Ashton and Cove, 1977) and 1% plant agar. Cultures were grown for 1 month at 22 °C under 16/8-h light/dark periods at a light intensity of 75 µmol m^-2^ s^-1^. Photosystem II (PSII) and I (PSI) functions were simultaneously recorded on cultures (pre-darkened for 30 min) using a Dua-PAM100 system (www.walz.com) during light exposure by red actinic light (278 µmol photons m^−2^ s^−1^) for 16 min and using 300 ms saturating red light pulses (10,000 µmol photons m^−2^ s^−1^). PSII function was assessed by measurement of maximum quantum yield of PSII photochemistry F_V_/F_M_ = (F_M_ − F_0_)/F_M_, where F_0_ and F_M_ represent a minimal and maximal Chl fluorescence, respectively, and non-photochemical quenching (NPQ) of maximal chlorophyll fluorescence under illumination (Bilger and Björkman 1990). The PSI quantum yields were further determined (Klughammer and Schreiber, 2008), including Y(I), quantum yield of non-photochemical energy dissipation due to PSI donor side limitation Y(ND) and quantum yield of non-photochemical energy dissipation due to PSI acceptor side limitation Y(NA).

Moss samples grown in the presence of 300 mM NaCl were lyophilized (20 mg each) for polyamine and amino acid determination (Ćavar Zeljković *et al*., 2024). The analysis was performed on a Nexera X2 UHPLC system coupled with an MS-8050 triple quadrupole mass spectrometer (www.shimadzu.com). Amino acids were separated chromatographically using an Acquity UPLC BEH AMIDE column (150 × 2.1 mm, 1.7 μm particle size) equipped with a matching pre-column. Polyamines, after derivatization with benzoyl chloride, were analyzed using an Acquity UPLC BEH C18 column (50 × 2.1 mm, 1.7 μm particle size) with a corresponding pre-column. Analyte identification was achieved in MRM mode by comparing the obtained spectra to those of authentic standards.

Samples for dicarboxylic and amino acid measurements were prepared from 200 mg of fresh material (FW), which was extracted with 1 ml of 100% methanol containing 0.1% formic acid. The supernatants were evaporated and redissolved in 20% methanol, then filtered on 0.2 μm centrifuge microfilters prior to UHPLC-ESI(-)-QTOF-MS (https://www.waters.com) using the ARION Polar C18 column (5 μm, 250 mm × 4.6 mm, https://www.chromservis.eu/) at 30 °C. Mobile phases A (acetonitrile) and B (10 mM formic acid) were mixed in gradient: 0 min 5% A, 0.1 min 5% A, 7 min 5% A, 12 min 35% A, 17 min 70% A, 17.5 min 100% A, 19 min 100% A, 19.5 min 5% A, 22 min 5% A. The mass spectrometer consisted of an electrospray ionization source operating in negative mode (ESI-) and a hybrid quadrupole time-of-flight (QTOF) mass analyzer (Synapt G2 Si, https://www.waters.com). The capillary and cone voltages were set to 2 kV and 15 kV, respectively. Source and desolvation temperatures were set at 120 °C and 500 °C. The collision energy was 20 eV. Centroided data were acquired in a data-dependent acquisition mode. The range of scanned masses was 70–1500 Da and 50–1500 Da. Quantitative analysis of metabolites was performed using external calibration. Oxidized glutathione was putatively identified by comparing the MS/MS product-ion spectrum of the detected metabolite with spectra from the NIST Tandem Mass Spectral Library (2023). The relative concentration was calculated from the peak areas (m/z 611) of the analyzed genotypes and corresponding controls (n = 6).

### RNA-seq data analysis

Moss WT and *aldh* mutants were grown on solid Knop medium with 300 mM NaCl, and the RNA was extracted using the RNeasy Plant Mini kit (www.quiagen.com). PolyA-enriched RNA was used for strand-specific cDNA library synthesis using the KAPA mRNA HyperPrep Kit (Roche). Sequencing on the NovaSeq X Plus instrument (www.illumina.com) at the Institute of Experimental Botany (https://ueb.cas.cz) resulted in at least 20 million 150-bp-long read pairs. Raw reads were quality-filtered using Rcorrector and Trim Galore scripts (Song and Florea, 2015). Levels of transcript expression (transcript abundances quantified as transcripts per million – TPM) were determined using Salmon (Patro *et al*. 2017) with parameters --posBias, --seqBias, --gcBias, --numBootstraps 30. The reference index was built from the *P. patens* CDS dataset v6.1 (Bi *et al*., 2024). Visualization, quality control of data analysis, and determination of differentially expressed genes were selected using the Sleuth (version 0.29.0) package in R software (Pimentel *et al*. 2017). Transcripts with *q*-value (*p*_adj_) ≤ 0.05 and log_2_ fold change ≥ 1 (upregulated) or ≤ -1 (downregulated) were considered to be significantly differentially expressed. Gene IDs from Phytozome (https://phytozome-next.jgi.doe.gov/) and Ensembl databases (https://www.ensembl.org) were used to assign identifiers such as Panther_ID (https://pantherdb.org/), PAC transcripts ID, EC number, KOG (http://eggnog6.embl.de/) and KEGG Orthology (KO) database (https://www.genome.jp/kegg/ko.html).

## RESULTS

### ALDH superfamilies in moss and barley and their relative expression

Whereas *ALDH* gene nomenclature in *Physcomitrium patens* has been published (Brocker *et al*., 2013), no information is available for *Hordeum vulgare*, one of the most important cereals worldwide. Therefore, we searched the barley genome (cv Morex v3, Mascher *et al*., 2021) and compared identified *ALDHs* with those in the moss genome. Both *ALDH* gene nomenclatures are listed in Table 1 and Supplementary Tables S6 and S7.

**Table 1.**
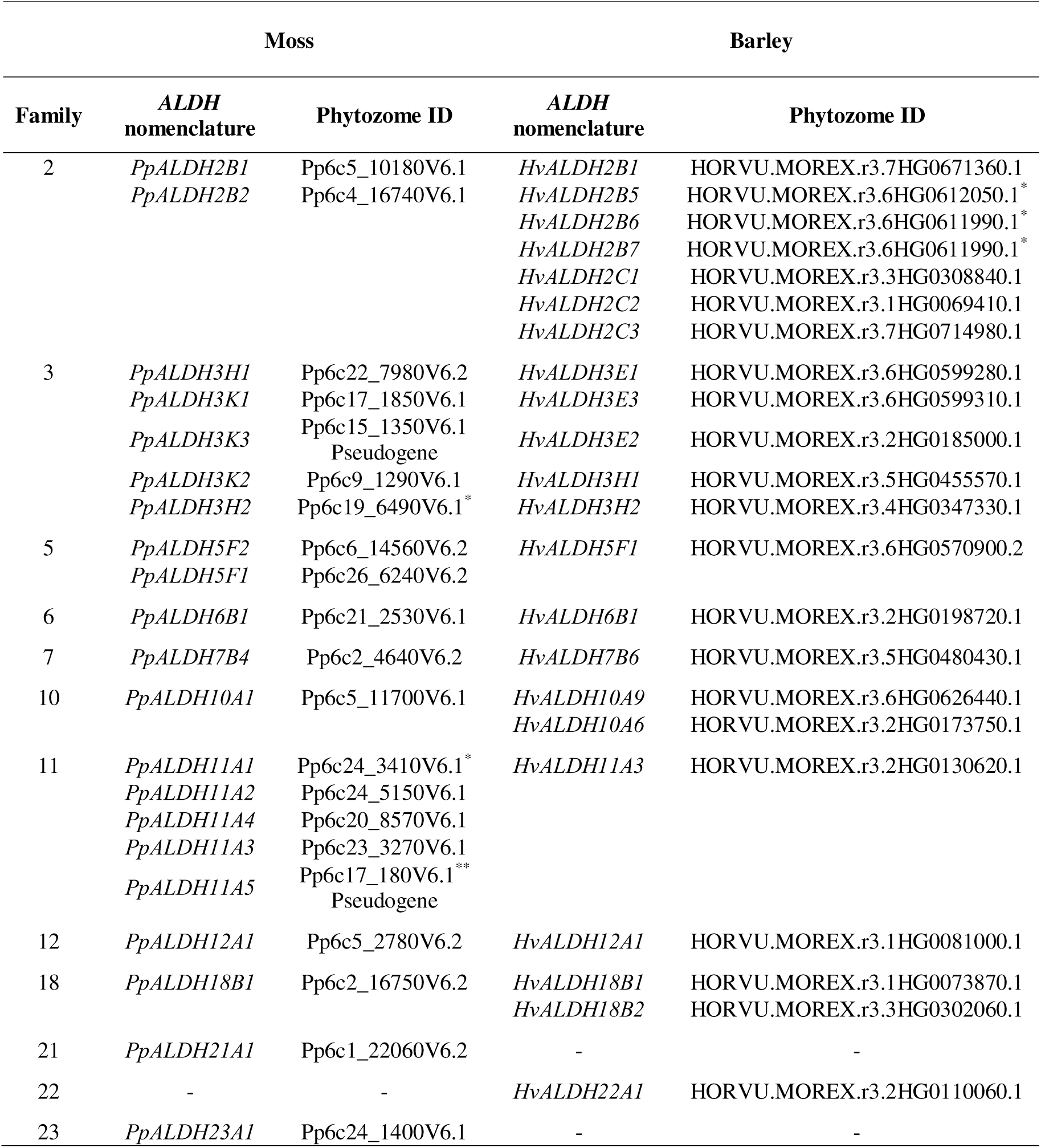
Gene nomenclature of *Physcomitrium patens* and *Hordeum vulgare ALDH* superfamilies. The genome of *Physcomitrium patens* v6.1 Gransden 2004 (Bi *et al*., 2024) and the genome of barley cv. MorexV3 (Mascher *et al*., 2021) were used for the *ALDH* sequence searches. Barley gene nomenclature follows the published closest monocot rice (Brocker *et al*., 2013). *Corrected sequence based on sequence alignments comprising the same family from other plants. **Presence of stop codon in exon 9.

While 19 genes and 2 pseudogenes are present in P. patens, 22 genes are found in the barley genome. Barley lacks *ALDH21* and *ALDH23* genes but possesses the *ALDH22* gene, which is absent in moss. In contrast to the four *ALDH11* genes and one *ALDH11* pseudogene (Supplementary Fig. S1) in moss, barley possesses only one *ALDH11* gene coding for a non-phosphorylating glyceraldehyde 3-phosphate dehydrogenase (GAPDH) known to regulate glycolysis as well as the synthesis of reduced sugars such as mannitol (Gao and Loescher, 2000; Rius *et al*., 2006). The ALDH2 family in barley comprises seven putative members compared to two in moss. The ALDH3 family contains five putative members in both lineages, although PpALDH3K3 is likely a pseudogene. Taken together, 12 out of 22 barley ALDHs should preferentially oxidize both aliphatic and aromatic aldehydes such as acetaldehyde, hexanal, nonanal, benzaldehyde, or cell wall-linked sinapaldehyde and coniferaldehyde as reported for other species (Nair *et al*., 2004; Long *et al*., 2009; Končitíková *et al*., 2015; Kirch *et al*., 2001; Sunkar *et al*., 2003; Demurtas *et al*., 2018). We found that three identical *HvALDH2B* gene copies with three promoters spanning a ∼ 41 kbp region of chromosome 6 were incorrectly annotated. The first gene copy, *HvALDH2B5* (annotated as HORVU.MOREX.r3.6HG0612050.1), lacks the first two exons. The second and third copies, *HvALDH2B6* and *HvALDH2B7*, are both under the same annotation as *HvALDH2B5* (Supplementary Fig. S1). When properly assembled, they all three encode three 543 aa-long ALDH2 enzymes (Supplementary Table S7). All assembled sequences are given in Supplementary dataset S1.

Moss genome contains 4 genes associated with the GABA shunt from the ALDH5, ALDH10 and ALDH21 families (Fig. 1): *PpALDH10A1*, two *ALDH5* genes (*PpALDH5F1* and *PpALDH5F2*) and *PpALDH21A1* (Brocker *et al*., 2013). The barley genome possesses only three *ALDH* genes linked to the GABA shunt: one *ALDH5* gene (*HvALDH5F1*) and two *ALDH10* genes (*HvALDH10A6* and *HvALDH10A9*). A qPCR analysis under standard conditions (Supplementary Fig. S1) using highly specific dual-labeled probes (listed in Supplementary Table S1 and Table S2) showed that transcripts of the GABA shunt-related *ALDH* genes listed above are highly abundant.

Overall, when comparing all *ALDH* genes in moss, *PpALDH3H1* and *PpALDH11A2* are poorly expressed, although both genes are complete. In barley, all *ALDH* genes are well expressed, except for the triplicated gene *HvALDH2B5/6/7*, whose transcript levels are very low and elevated only under specific stress or hormone treatments.

### Stress-responsive ALDHs in moss and barley

Gene expression is regulated by *cis*-regulatory elements, promoter and enhancer architecture, silencers and others (Vihervaara *et al*., 2017). The promoter analysis using the Plant-CARE database revealed a high number of stress response-related elements in moss and barley *ALDH* superfamilies (Supplementary Fig. S2), including drought-responsive MYB sites, stress-responsive (STRE) and ABA-responsive elements (ABRE) known to be linked to drought stress adaptation (Wang *et al*., 2025). The presence of a motif does not solely guarantee a change in expression due to varying chromatin accessibility, binding site occupancy, or other factors. Therefore, we studied by qPCR changes in expression upon various abiotic stresses (drought, salt, nitrogen, heat and cold) and several hormone treatments including ABA, MeJA, auxin and cytokinin (Fig. 2).

**Fig. 2.**
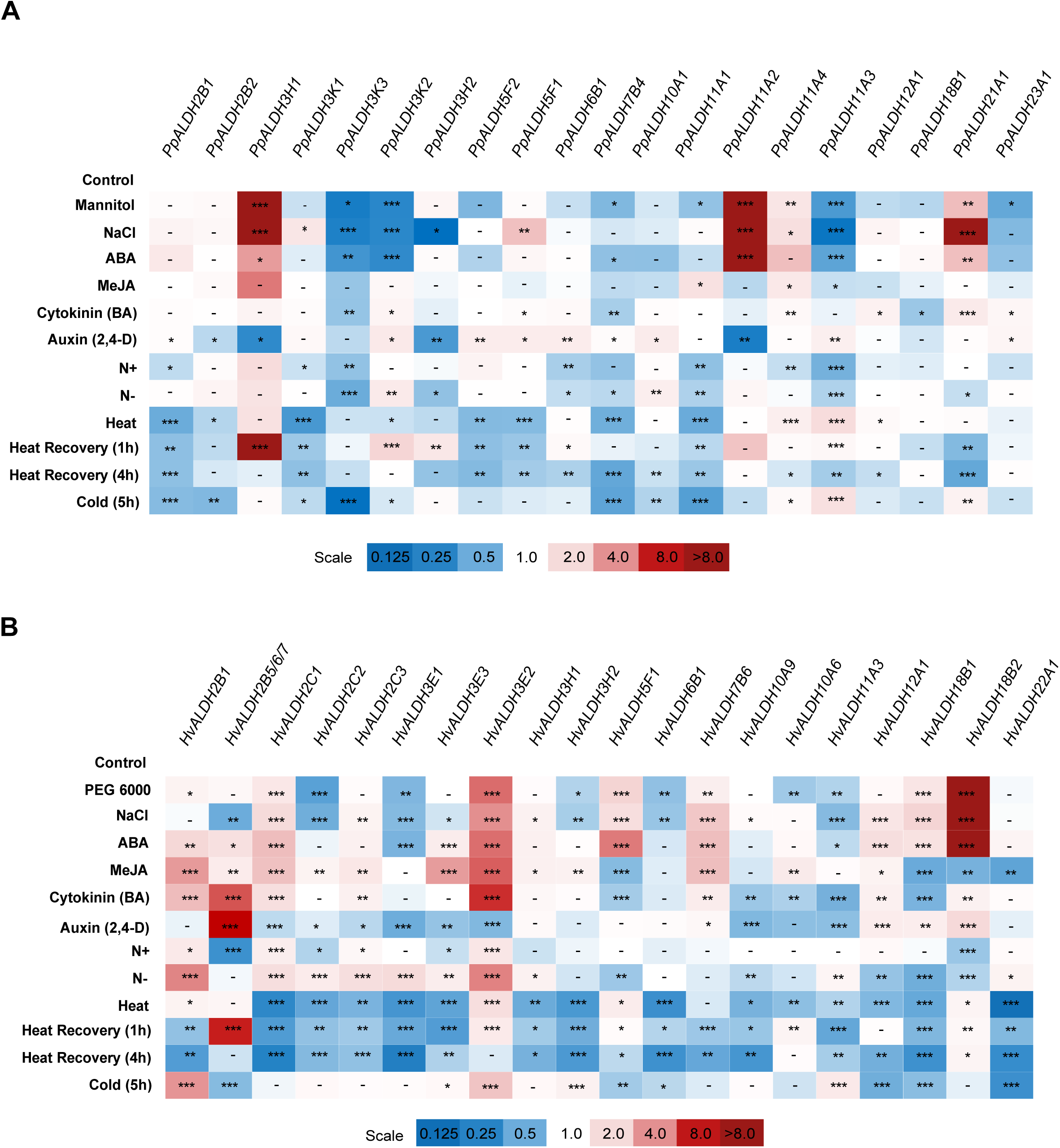
Gene expression screening in moss and barley seedlings upon hormone or stress treatments. **(A)** Expression profiles of moss *ALDH* superfamily. **(B)** Expression profiles of barley *ALDH* superfamily. Moss was grown in Knop’s medium, barley was grown in Hoagland’s solution and expression was determined by qPCR using dual-labeled FAM-TAM probes. Fold change in relative expression vs control in six technical replicates was statistically analyzed with a t-test; *, **, and *** correspond to p-values of 0.05 > p > 0.01, 0.01 > p > 0.001, and p < 0.001, respectively.

In moss, salt and osmotic stresses, as well as ABA treatment, resulted in a strong upregulation of four genes from the ALDH3, ALDH11 and ALDH21 families (*PpALDH3H1*, *PpALDH11A2*, *PpALDH11A4* and *PpALDH21A1*), but also in a strong downregulation of three genes from the same families: *PpALDH3K2, PpALDH3K3*, and *PpALDH11A3* (Fig. 2A). Heat and cold stresses generally decreased the expression of *PpALDH* genes, except for *PpALDH11A3* and *PpALDH11A4*, which were upregulated. During a 1-hour heat recovery period, the three *PpALDH3* genes, *PpALDH3H1, PpALDH3H2* and *PpALDH3K2*, were upregulated (Fig. 2A). Notably, the expression of either *PpALDH12* or *PpALDH18*, which are associated with the regulation of proline metabolism, was not significantly altered. Responses to MeJA were mild and not statistically significant. Cytokinin treatment resulted in the upregulation of *PpALDH21A1* along with *PpALDH11A4* and the downregulation of both *PpALDH3K3* and *PpALDH7B4*. The responses to auxin treatments were stronger, with several genes mildly upregulated, including those from families 5 and 10 and three genes (*PpALDH3H1*, *PpALDH3H2*, and *PpALDH11A2*) downregulated.

In barley, stress responses were very different from moss within the same ALDH families. Salt and osmotic stresses, as well as ABA treatment, resulted in a strong upregulation of seven genes from six different families: ALDH2, ALDH3, ALDH5, ALDH7, ALDH12, and ALDH18. In contrast, moss showed upregulation of only three genes, all from the ALDH3 family. The barley genes from *ALDH12 and ALDH18* families are involved in proline metabolism, those from *ALDH2* and *ALDH3* are linked to cell wall lignification and lipid peroxidation, barley *ALDH7B6* gene contributes to lysine catabolism and the *HvALDH5F1* gene is linked to the GABA shunt (Fig. 2B). On the contrary, the unique *ALDH11* gene (*HvALDH11A3*) was downregulated as well as *HvALDH2C2* and *HvALDH3E1.* Similar to moss, heat and cold stresses generally decreased the expression of most barley *ALDH* genes (Fig. 2B). The *HvALDH3E2* gene was upregulated by heat stress, and three genes, *HvALDH2B5, HvALDH3E2* and *HvALDH10A9,* accumulated during heat recovery (Fig. 2B). Cold stress induced the expression of four genes, namely *HvALDH11A3*, *HvALDH2B1*, *HvALDH3E2* and *HvALDH3H2.* Hormone responses were more pronounced in barley compared to moss. MeJA induced the expression of many *HvALDH*2 and *HvALDH3* genes, plus *HvALDH7B6* and strongly downregulated the expression of *HvALDH5F1*, *HvALDH18B1/B2* and *HvALDH22A1* genes. Cytokinin also upregulated several *HvALDH2* and *HvALDH3* genes and downregulated *ALDH* genes from families 5 and 10. Auxin upregulated *HvALDH12A1*, *HvALDH18B1* and *HvALDH18B2*, plus *HvALDH2B5.* While the excess or lack of nitrogen did not induce strong responses in moss, several genes from the *H*v*ALDH2* and *HvALDH3* families were highly upregulated.

Regarding the GABA shunt-associated *ALDHs* in moss, the expression of both *ALDH5* genes decreased upon exposure to heat stress and upon heat recovery. Transcript levels of *PpALDH5F1* increased only upon salt stress. Transcripts of *PpALDH21A1* increased upon drought, salt, and ABA exposure, whilst a mild elevation was observed during cold stress and a decline during the heat recovery period. Expression of *PpALDH10A1* increased only upon nitrogen deprivation. In barley, the sole *HvALDH5F1* gene was highly upregulated by drought, salt and ABA and slightly less by heat stress. The two *HvALDH10A6* and *HvALDH10A9* were primarily downregulated in response to various treatments.

### Moss ALDH10A1 oxidizes ***ω***-aminoaldehydes to produce ***β***-alanine and GABA

Focusing on the GABA shunt, we cloned and expressed Δ*_1-52_PpALDH10A1* (507 aa), comprising the SKL peroxisomal signal at its C-terminus found in the majority of ALDH10 members. The enzyme is dimeric in solution as verified by gel permeation chromatography (Supplementary Fig. S3) in line with reported observations on the ALDH10 family (Tylichová *et al*., 2010; Díaz-Sánchez *et al*., 2012; Kopečný *et al*., 2013). The stability of the enzyme rose by ∼ 5 °C in the presence of the coenzyme.

From numerous aldehydes screened as potential substrates for PpALDH10A1 using NAD^+^ as the coenzyme, ω-aminoaldehydes such as APAL, ABAL, GBAL and TMABAL were identified as preferred substrates (Fig. 3A). The enzyme exhibited the highest specific activities with APAL, followed by its acetylated form, acAPAL, which occurs naturally during polyamine oxidation of Spermine (Spm), Spermidine (Spd), *N*^1^-acSpm or *N*^1^-acSpd. Kinetic properties of PpALDH10A1 with its preferred substrates were explored further (Fig. 3B and Table 2). The enzyme showed the highest catalytic efficiency (*k*_cat/_*K*_m_) values with GBAL (7.6 × 10^5^ s^−1^M^−1^) and APAL (7.2 × 10^5^ s^−1^M^−1^), followed by acAPAL and TMABAL. The highest observed *k*_cat_ values reached 6.5 s^−1^ and *K*_m_ values were in the low micromolar range. The apparent catalytic efficiency value for NAD^+^ was 1600-fold higher than for NADP^+^, when using 0.1 mM APAL, due to the high *K*_m_ value of 32 mM for NADP^+^ versus 40 µM for NAD^+^. PpALDH10A1 exhibited substrate inhibition kinetics with various aminoaldehydes.

**Fig. 3.**
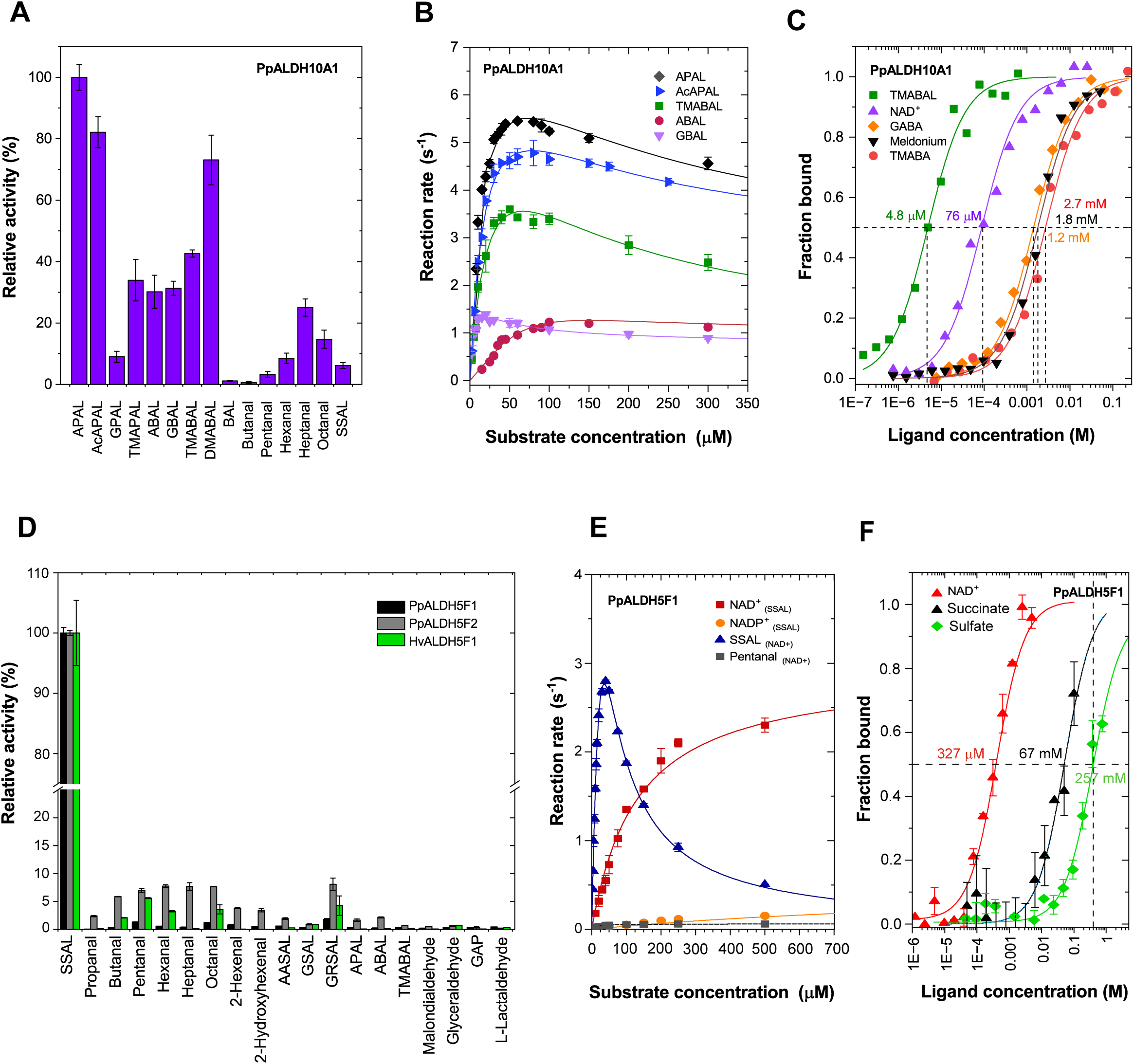
Kinetic properties of ΔPpALDH10A1 and PpALDH5 family. **(A)** Substrate screening for major substrates of Δ PpALDH10A1. Measured with 1 mM substrate in 150 mM Tris-HCl buffer (pH 8.5) containing 1 mM NAD^+^ at 30 °C. Error bars stand for S.D. (n=4). **(B)** Saturation curves for five aminoaldehydes. Measured in 150 mM Tris-HCl buffer (pH 8.5) containing 1 mM NAD^+^ at 30 °C. **(C)** Affinity measurements of ΔPpALDH10A1 for TMABAL, NAD^+^, reaction products TMABA and GABA and the product analog meldonium by MST on a Monolith NT.115 instrument (NanoTem-per Technologies) at 25 °C. The binding was assessed in 50 mM Tris-HCl at pH 9, 100 mM NaCl, and 1% GOL, supple-mented with 0.05% Tween. *K*_D_ values are indicated. **(D)** Screening of substrate specificity of moss and barley ALDH5. Measured with 200 µM aldehyde in 150 mM HEPES buffer (pH 8.0) and 1.5 mM NAD^+^ at 30 °C. **(E)** Saturation curves for SSAL, NAD^+^, NADP^+^ and pentanal with PpALDH5F1. Data were measured in 150 mM HEPES buffer pH 8.0, using 1.5 mM NAD^+^ (for SSAL) or 40 μM SSAL (for coenzymes). Error bars stand for S.D. (n=4). **(F)** Affinity measurements of PpALDH5F1 for NAD^+^, succinate and sulfate ions by MST on a Monolith X instrument (NanoTemper Technologies) at 25 °C. The binding was assessed in 50 mM Tris-HCl at pH 8.0, 100 mM NaCl, supplemented with 0.05% Tween. *K*_D_ values are indicated.

**Table 2.**
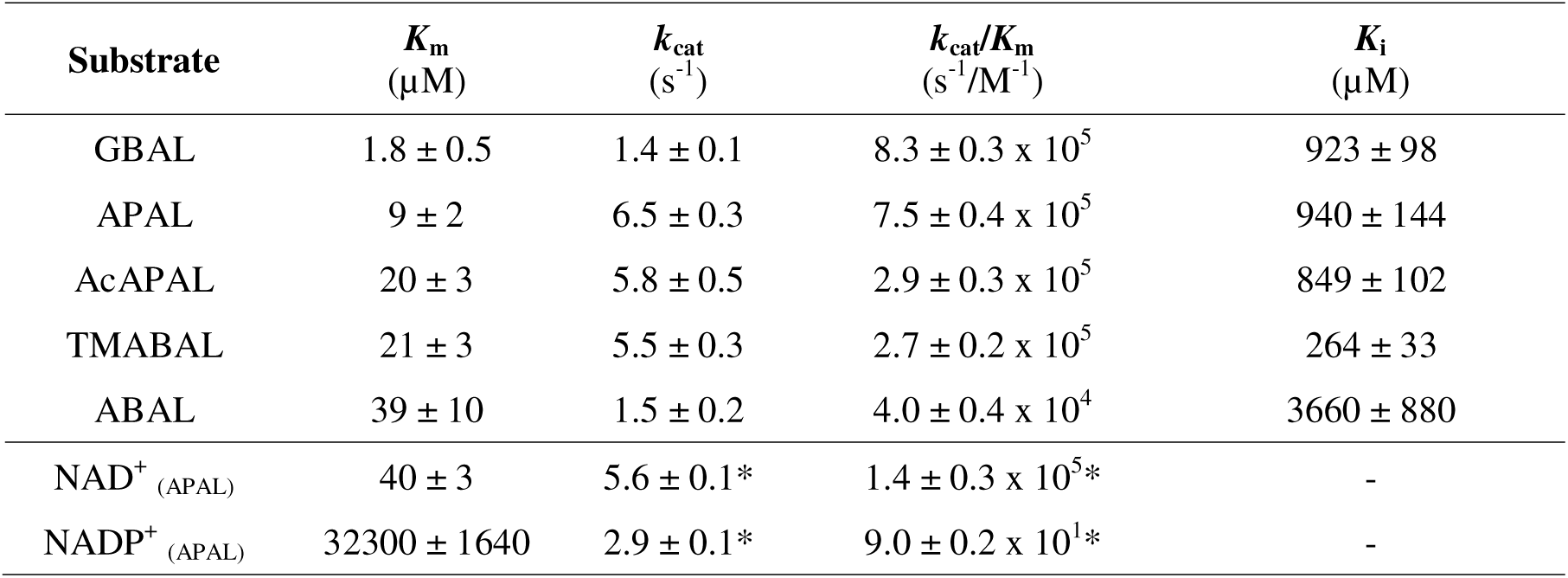
Kinetic parameters for major substrates of. Δ**PpALDH10A1.** Saturation curves were measured in 150 mM Tris-HCl buffer, pH 8.5, and 1 mM NAD^+^ at 30 °C. Saturation curves for NAD(P)^+^ were measured with 0.1 mM APAL, giving the highest rates. The lower *k*_cat_ values for coenzyme (indicated by asterisks), compared to those for APAL, result from a fixed sub-saturating APAL concentration used for the NAD^+^ saturation curve measurement. Kinetic constants, including their S.E. values, were determined using GraphPad Prism 8.0 software.

We verified the affinity to substrate and product by microscale thermophoresis (Fig. 3C). As expected, affinity values for the reaction products GABA and TMABA, as well as for the product analog meldonium, were in the low millimolar range in contrast to low micromolar concentration measured for TMABAL (Fig. 3C). Betaine aldehyde (BAL) was a poor substrate, due to the presence of an isoleucine residue in the active site, which has been shown to impair BAL binding and oxidation in both spinach and maize ALDH10 (Díaz-Sánchez *et al*., 2012; Kopečný *et al*., 2013). Substrate specificity of two barley ALDH10 enzymes was previously analyzed with APAL, ABAL, TMABAL and BAL (Fujiwara *et al*., 2008), and thus they were not analyzed in our work.

### Moss and barley ALDH5 oxidize SSAL to produce succinate

PpALDH5F1 (492 aa), PpALDH5F2 (498 aa) and HvALDH5F1 (526 aa) were produced in *E. coli* as tetramers of ∼ 230 kDa, as observed for human ALDH5A1 (Kim *et al*., 2009). PpALDH5F1 and PpALDH5F2 share 76% sequence identity as well as 69% and 65% sequence identity with HvALDHF1, respectively. All three enzymes displayed a strong substrate preference for SSAL (Fig. 3D), whereas other aliphatic aldehydes, ω-aminoaldehydes, α-aminoadipate-semialdehyde (AASAL, major substrate of ALDH7), Glu γ-semialdehyde (GSAL, major substrate of ALDH12), L-lactaldehyde and GAP (the major substrate of ALDH11) were all very poor substrates. Only PpALDH5F2 showed a low activity with several aliphatic aldehydes and glutaric semialdehyde (GRSAL) (up to ∼ 5% of SSAL activity).

The kinetic properties measured with SSAL (Table 3) showed that PpALDH5F1 displayed a 5-times higher catalytic efficiency than PpALDH5F2 and a similar to HvALDH5F1. *K*_m_ values for SSAL ranged from ∼ 60 to 90 μM concentration. All three ALDH5 enzymes are more active with NAD^+^ compared to NADP^+^ and a substrate inhibition effect was observed at high SSAL concentrations. *K*_m_ values of ∼ 100 µM were measured for NAD^+^, while *K*_m_ values for NADP^+^ were in the range of 1.5 - 2.0 mM (Fig. 3E). Stability of PpALDH5F1 lowered by ∼ 4 °C upon added coenzyme as measured by nano-DSF and the affinity to NAD^+^ and the succinate (reaction product) to PpALDH5F1 was found ∼ 327 µM and 67 mM, respectively, as found by MST (Fig. 3F).

**Table 3.**
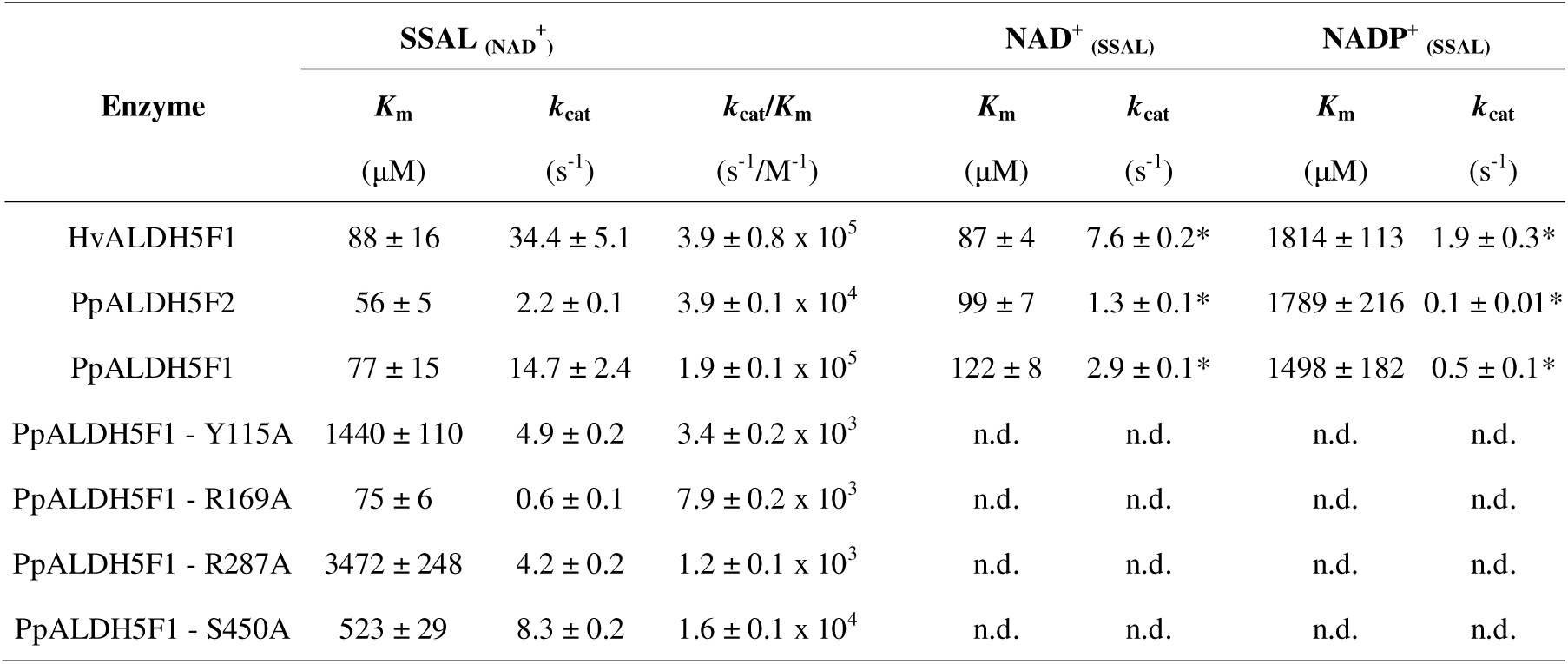
Kinetic parameters for the major substrate SSAL and the NAD^+^ coenzyme of plant ALDH5 variants. Saturation curves for SSAL were measured in 150 mM HEPES buffer, pH 8.0, using 1.5 mM NAD^+^ at 30 °C; those for NAD^+^ were measured using SSAL concentration providing the highest rates, i.e., 200 μM for PpALDH5F2 and 40 μM for PpALDH5F1/HvALDH5F1. The lower *k*_cat_ values for NAD^+^ (indicated by asterisks) compared to those for SSAL result from using a fixed sub-saturating SSAL concentration due to substrate inhibition effect at saturating concentration. Kinetic constants, including their S.E. values, were determined using GraphPad Prism 8.0 software. n.d. stands for not determined

| Enzyme | SSAL (NAD<sup>+</sup>) |  |  | NAD<sup>+</sup> (SSAL) |  | NADP<sup>+</sup> (SSAL) |  |
| --- | --- | --- | --- | --- | --- | --- | --- |
| | $K_m$ | $k_{\text{cat}}$ | $k_{\text{cat}}/K_m$ | $K_m$ | $k_{\text{cat}}$ | $K_m$ | $k_{\text{cat}}$ |
| | (μM) | (s<sup>-1</sup>) | (s<sup>-1</sup>/M<sup>-1</sup>) | (μM) | (s<sup>-1</sup>) | (μM) | (s<sup>-1</sup>) |
| HvALDH5F1 | 88 ± 16 | 34.4 ± 5.1 | 3.9 ± 0.8 $\times 10^5$ | 87 ± 4 | 7.6 ± 0.2* | 1814 ± 113 | 1.9 ± 0.3* |
| PpALDH5F2 | 56 ± 5 | 2.2 ± 0.1 | 3.9 ± 0.1 $\times 10^4$ | 99 ± 7 | 1.3 ± 0.1* | 1789 ± 216 | 0.1 ± 0.01* |
| PpALDH5F1 | 77 ± 15 | 14.7 ± 2.4 | 1.9 ± 0.1 $\times 10^5$ | 122 ± 8 | 2.9 ± 0.1* | 1498 ± 182 | 0.5 ± 0.1* |
| PpALDH5F1 - Y115A | 1440 ± 110 | 4.9 ± 0.2 | 3.4 ± 0.2 $\times 10^3$ | n.d. | n.d. | n.d. | n.d. |
| PpALDH5F1 - R169A | 75 ± 6 | 0.6 ± 0.1 | 7.9 ± 0.2 $\times 10^3$ | n.d. | n.d. | n.d. | n.d. |
| PpALDH5F1 - R287A | 3472 ± 248 | 4.2 ± 0.2 | 1.2 ± 0.1 $\times 10^3$ | n.d. | n.d. | n.d. | n.d. |
| PpALDH5F1 - S450A | 523 ± 29 | 8.3 ± 0.2 | 1.6 ± 0.1 $\times 10^4$ | n.d. | n.d. | n.d. | n.d. |

### Plant ALDH5 is a tetramer and the coenzyme adopts two conformational states

Since the dimeric structures of several plant ALDH10 members were published (Tylichová *et al*., 2010; Díaz-Sánchez *et al*., 2012; Kopečný *et al*., 2013) as well as the tetrameric structure of ALDH21 (Kopečná *et al*., 2017), here, we focused on the first structure of plant ALDH5, namely PpALDH5F1. The apoform structure (PDB 8OF3), solved at 1.35 Å resolution, contains two monomers in its asymmetric unit, which form the known “dimer-of-dimers” tetramer by crystal symmetry (Figure 4A) in line with the gel permeation chromatography (Supplementary Fig. S3). The overall fold of plant ALDH5 (Fig. 4A) resembles that of other ALDH superfamily members, including *E. coli* NADP^+^-dependent SSALDH (PDB 3JZ4, Langendorf *et al*., 2010), mitochondrial human ALDH5A1 (PDB 2W8Q, Kim *et al*., 2009), plant ALDH2 (PDB 4PXL and 4PZ2), ALDH7 (PDB 4PXN, Končitiková *et al*., 2015), ALDH12 (PDB 6D97, Korasick *et al*., 2019) and ALDH21 (PDB 5MZ8, Kopečná *et al*., 2017). Two PpALDH5F1 monomers are arranged as a domain-swapped dimer. Each monomer consists of three domains (Fig. 4B): an NAD^+^-binding fold (residues 1–130, 152–260, and 468–476), a catalytic α/β fold (residues 261–448), and a C-terminal oligomerization flap (residues 131–151 and 477–492). The inter-domain linker (residues 449–468) forms a lid on the active site and creates an anchor loop that stabilizes the aldehyde substrate. The catalytic Cys293 is located on a loop that spans the crevice between the NAD^+^-binding and catalytic domains.

**Fig. 4.**
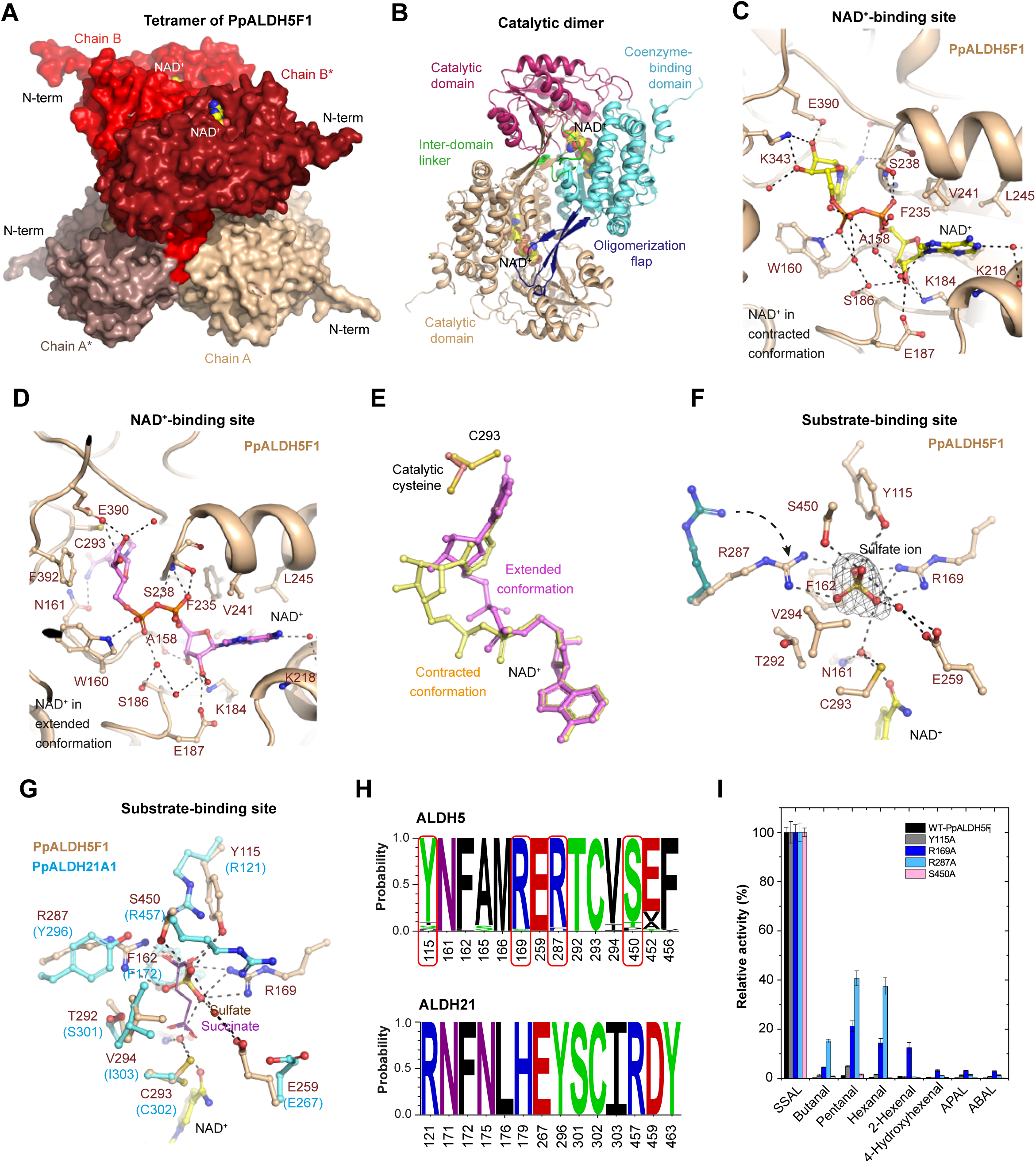
Structure of PpALDH5F1 and residues linked to SSAL binding. **(A)** The tetrameric structure of PpALDH5F1 is composed of four monomers arranged as two domain-swapped dimers. **(B)** Each subunit contains catalytic (red), coenzyme-binding (aquamarine), and oligomerization (dark blue) domains. The NAD^+^ coenzyme is in yellow. **(C)** NAD^+^ molecule bound in the contracted conformation. Interacting residues are labeled and the coenzyme is in yellow. **(D)** NAD^+^ molecule bound in the contracted conformation and the coenzy-me is colored in violet. **(E)** Superposition of the coenzyme in extended (in violet) and contracted (in yellow) conformations, next to the catalytic cysteine. **(F)** Active-site residues forming the substrate channel in PpALDH5F1. The bound sulfate mimicking the substrate is shown in its annealing Fo-Fc omit map (black mesh) contoured at 3.0 σ. **(G)** Superposition of the substrate channels of PpALDH5F1 (light brown, this work) and PpALDH21 (cyan, PDB 5MZ8, Kopečná *et al*., 2017). The succinate in purple line representation bound to the active site of PpALDH21 is shown for comparison. **(H)** Conservation of amino acids forming the substrate channel in ALDH5 and ALDH21 families. Residue numbering follows that of PpALDH5F1 and PpALDH21. Residues binding the carboxylate group of SSAL are highlighted in red rectangles. Sequence logos were made using WebLogo 3 (http://weblogo.threeplusone.com). **(I)** Substrate specificity of PpALDH5F1 variants. Measured with 200 µM aldehydes in 150 mM HEPES buffer (pH 8.0) at 30 °C. Error bars stand for S.D. (n=4).

We also solved two NAD^+^-PpALDH5F1 structures (PDB 8OF1 and 8OFM, Supplementary Table S4) isomorphous to the apoform structure, indicating that the bound coenzyme did not induce large changes. In the PDB 8OF1, the NAD^+^ adopts a contracted conformation in both subunits, while in the PDB 8OFM, the NAD^+^ has an extended conformation in both subunits (Fig. 4C, 4D and 4E). While the extended conformation of NAD^+^ is more frequent, the contracted conformation is more typical of NADH (Hammen *et al*., 2002). The adenosine moiety of NAD^+^ lies in a hydrophobic pocket flanked by two helices (residues 197-206 and 218-232) and the ribose is bound by multiple H-bonds to the side chains of Lys184 and Glu187 and the main-chain O atom of Ala158. Two α-phosphate oxygens, O1A and O3, bind to the main-chain NH atom and the side-chain OG atom of Ser238, while the oxygen atom of the β-phosphate group interacts with the NE1 atom of Trp160 (Fig. 4C and 4D). In the contracted conformation, the nicotinamide ribose establishes a bidentate H-bond with a conserved lysine residue (Lys343), whereas in the extended form, the ribose moves and establishes a bidentate H-bond with a conserved Glu residue (Glu390).

### Binding of SSAL requires a tyrosine and two arginine residues

Our attempts to obtain a PpALDH5F1 structure with the reaction product succinate bound to the active site, either by co-crystallization or soaking experiments, were unsuccessful. The primary issue was related to the crystallization condition comprising 1.7 M lithium sulfate, in which 100 mM succinate did not outcompete the observed sulfate ions occupying the active sites in both NAD^+^ structures (Fig. 4). Binding of sulfate ion at high millimolar concentrations was confirmed by MST (Fig. 3F)

Sulfate binds at the position where the carboxylate group of SSAL/succinate would be located (Fig. 4G) and interacts with two basic residues, Arg169 and Arg287, as well as with Ser450 and Tyr115, all located in the upper part of the substrate channel. The side chain of Arg287 is flexible and adopts different conformations (Fig. 4F). The other conserved and nonpolar residues, which maintain the active site geometry, include Phe162, Ala165, Met166, Thr292, Val294 and Phe456. The catalytic cysteine Cys293, the general base Glu259 and the oxyanion hole residue Asn161 are located in the lower part of the channel. Asn161, which donates a hydrogen bond to the O atom of the aldehyde group of SSAL, is positioned on a loop adjacent to the catalytic loop. All interacting and surrounding residues are highly conserved within the ALDH5 family (Fig. 4H) but differ markedly from those in the ALDH21 family, even though both families catalyze the same reaction.

Four key residues for SSAL oxidation have been studied by site-directed mutagenesis (Table 3 and Fig. 4I) and the sequence alignment including the considered mutations is shown in Supplementary Fig. S4. Both Arg169 and Arg287 are essential for catalysis, as the R169A and R287A variants displayed low catalytic efficiencies (*k*_cat_/*K*_m_ ratios) for SSAL corresponding to 3.9 and 0.6% of the WT values. While the R169A variant is affected in *k*_cat_ value (25-fold lower compared to WT), the R287A variant is strongly affected in *K*_m_ value (45-fold higher compared to WT). The catalytic efficiencies of Y115A and S450A variants were reduced to ∼ 1.8 and 8.0% of the WT. Thus, the Y115 residue is also essential for binding, as the *K*_m_ value of the Y115A mutant is 19-fold higher compared to WT. The removal of either Arg residue reduced the specificity towards negatively charged aldehyde and increased the relative oxidation of aliphatic aldehydes such as pentanal or hexanal, but was not sufficient to increase the oxidation of aminoaldehydes, major substrates of the ALDH10 family (Fig. 4I).

### Slower growth of aldh knockouts

*P. patens* is highly tolerant to abiotic stresses, including salt stress up to 350 mM NaCl concentration and osmotic stress induced by sorbitol up to 500 mM concentration (Frank *et al*., 2005). We analyzed the function of three genes *in vivo*, namely *PpALDH5F2*, *PpALDH10A1* and *PpALDH21A1*, all involved in the GABA shunt pathway, by generating knockout lines by homologous recombination (Supplementary Fig. S5). Notably, we were unable to generate *aldh5F1* knockout lines. Three independent lines for each knockout were grown in liquid Knop medium for 1 month and then transferred to 6-well plates containing modified Knop medium with agar only or in the presence of mannitol (up to 1000 mOsm/kg) and sodium chloride (up to 300 mM). Colony diameter was measured in time (Fig. 5B and 5C) as well as the number of early gametophores (Fig. 5A). Under normal conditions, as well as under both osmotic and salt stress, the growth of *aldh5F2* knockout lines was slower than that of the wild type (WT), with colony size reduced to approximately 75% of WT. Both *aldh10A1* and *aldh21A1* knockouts were less affected, although they also exhibited slower growth. The number of early gametophores was slightly higher in all three knockouts, suggesting active development; however, the differences between independent lines for each knockout were substantial, resulting in high variability.

**Fig. 5.**
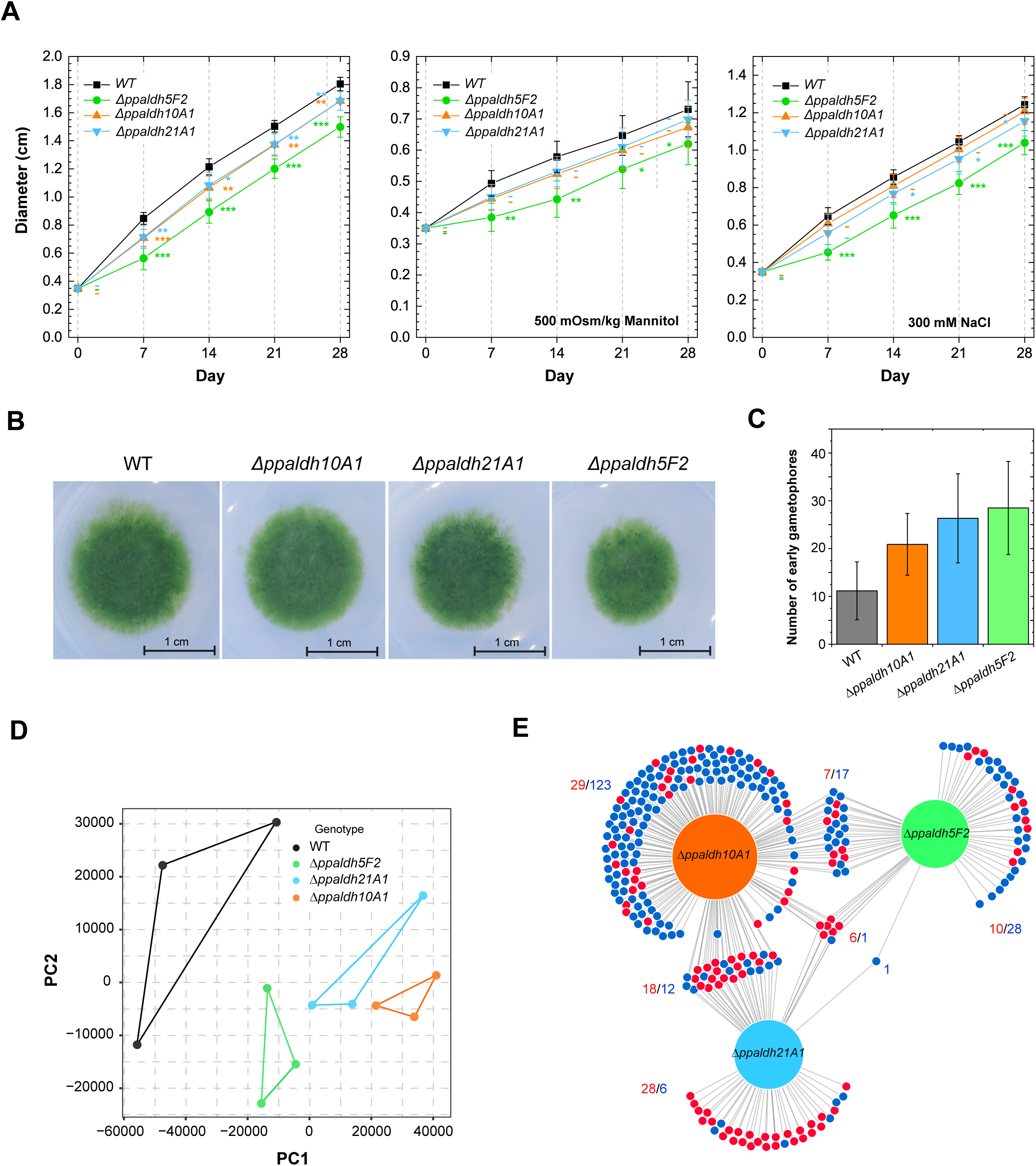
Growth differences and RNA-seq in moss *aldh* knockouts. **(A)** Growth after point inoculation on modified Knop medium and 1% plant agar. Cultivated for one month under normal, osmotic stress and salt stress conditions at 19 °C. Graph values indicate mean ± S.D. (n=15). **(B)** Phenotype of moss *aldh* knockouts. **(C)** The number of early gametophores at 21 days after inoculation measured in a 3.8 mm^2^ area (n=10). **(D)** Principal Component Analysis of processed RNA-seq data generated by the Sleuth R package (v 0.30.1). PC1 accounts for the maximized variance, while PC2 is orthogonal to PC1 with the second-highest variance. **(E)** DiVenn diagrams depicting different and overlapping gene regulation in *aldh5F2*, *aldh10A1* and *aldh21A1* mutants in comparison with the wild-type. Red and blue dots represent significantly upregulated and downregulated genes (q ≤ 0.05, fold-change ≥ 2), respectively, in the linked comparisons.

### Transcriptome analyses revealed upregulated glutathione-S-transferase genes in aldh knockouts

We performed whole-transcriptome analysis of all three *aldh* knockouts and WT. Principal component analysis (PCA) was used for quality control assessment of the RNA-seq data (Fig. 5D) and three biological replicates per genotype were ultimately selected for determining differentially expressed genes (DEGs). Gene transcripts with *q*-value (*p*_adj_) ≤ 0.05 versus WT and log_2_-fold change greater than one were considered significantly upregulated, while those with log_2_-fold change lower than -1 were considered significantly downregulated (Supplementary Fig. S6) and they are all listed in the Supplementary Dataset S2. In the *aldh5* knockout, we identified 70 DEGs (23/47 genes up/downregulated), in the *aldh21* knockout, we found 72 DEGs (52/20 genes up/downregulated), and finally, in the aldh10 knockout, we found 213 DEGs (60/153 genes up/downregulated) (Fig. 5E). Their transcription levels in three biological replicates per genotype are shown as heat maps in Supplementary Fig. S7, S8 and S9.

Interestingly, transcription of 6 genes was strongly induced in all three knockouts, namely two *glutathione-S-tranferase* (*GST*) genes (Pp6c15_12200 and Pp6c23_10140), gene of *methylthioadenosine (MTA) nucleosidase* (*MTN*) (Pp6c26_3100), *cytoskeleton-associated protein 5 (CKAP5)* gene (Pp6c7_1120), *DDE superfamily endonuclease* gene (Pp6c15_4090) and *disease resistance protein/protein kinase gene* (Pp6c20_10940). MTN hydrolyzes MTA, a by-product of SAM-dependent methylation reactions, such as polyamine biosynthesis, to methylthioribose, which continues into the Met recycling pathway. The only common downregulated gene codes for ubiquitin-conjugating enzyme E2Q (Pp6c23_10470), which participates in cell cycle regulation (Kraft *et al*., 2005).

Additional *GST* genes were upregulated in moss *aldh* mutants: transcripts of Pp6c1_17240, Pp6c4_6710 and Pp6c11_8010 were increased by ∼ 1.3- to 1.8-fold, transcripts of Pp6c12_990 were 1.4-fold higher, and the expression of Pp6c7_7560 was nearly two-fold higher than in WT. Above-encoded GSTs catalyze the conjugation of glutathione to various electrophiles and can contribute to cell protection. Consequently, we measured oxidized glutathione (GSSG) levels, which were reduced to 67 ± 15, 62 ± 13 and 50 ± 11% in *aldh5*, *aldh10* and *aldh21* knockout, respectively, compared to the WT.

Among other genes that had nearly 2-fold expression in all three *aldh* knockouts were Pp6c2_7060 and Pp6c1_8690. The first gene codes for a “dessication-induced Vicinal Oxygen Chelate 1 (VOC1)/glyoxalase-like protein”, but it is not a functional glyoxalase I/II, known to detoxify methylglyoxal to lactate using glutathione as a co-substrate and the significance of glyoxalase-like proteins is unclear (Schmitz *et al*., 2017). The latter gene encodes the RNA helicase DDX20 (Gemin3), known to be involved in transcription, post-transcriptional modifications, and mRNA splicing. Both *aldh21* and aldh10 knockouts have upregulated a *NatB* gene (Pp6c22_6900) encoding the N-terminal acetyltransferase linked to salt and osmotic stress responses (Huber *et al*., 2020). The other upregulated gene (Pp6c6_3070) in *aldh21* and *aldh10* knockouts encodes dioxygenase, annotated as “gibberellin A44 oxidase”. However, since moss produces only a 3β-hydroxy-kaurenoic acid derivative (Miyazaki *et al*., 2018) and not active gibberellins, its role in the metabolism of kaurenoic acid remains unclear.

All three knockouts also show reduced expression of fatty acid biosynthesis genes, including *3-oxoacyl acyl-carrier-protein synthase* (Pp6c12_3310), *omega-3 desaturase* (Pp6c26_6300) and *omega-6 desaturase* (Pp6c16_3180) genes. Both *aldh5* and *aldh10* have downregulated several genes coding for DNA-binding proteins or transcription factors, such as *dehydration-responsive element-binding protein 3* (Pp6c3_16580), *C2H2-type protein* (Pp6c11_1300) or *transcription factor CRF3* (Pp6c11_12740).

Finally, the gene (Pp6c21_2900) that is downregulated in *aldh5* and *aldh21* (to ∼45% of WT) and in *aldh10* (to 62%), encodes cytochrome *b_6_*, a core component of the cytochrome *b6f* complex involved in electron transport between PSII and PSI. Direct analysis of the redox state of PSI in all three knockouts revealed very similar profiles of quantum yield of PSI photochemistry Y(I), Y(ND) and Y(NA) (Supplementary Fig. S10), indicating that the reduced transcription of *cyt b_6_* does not limit the function of PSI. However, non-photochemical quenching (NPQ), a photoprotective mechanism that helps to prevent photodamage of PSII, was consistently altered in all three knockout lines compared to the WT. Finally, the F_V_/F_M_ ratio, a measure of PSII photosynthetic efficiency, was 0.86 ± 0.01 for WT and those for all three *aldh* knockouts were ∼ 0.84 (Supplementary Fig. S10), indicating that there was no significant disruption of PSII photochemistry.

### Metabolite changes in three aldh knockouts and expression of associated genes

The *aldh5F2* or *aldh21A1* knockouts accumulated approximately twice the amount of GABA compared to WT plants (Fig. 6A and Supplementary Table S9). Succinate levels were affected only in the *aldh21A1* knockout, where they were reduced by ∼ 20%, indicating that the lack of cytosolic SSALDH is harder to compensate for. Because GABA is synthesized from Glu by GAD, also Glu levels were approximately twice as high in both knockouts.

**Fig. 6.**
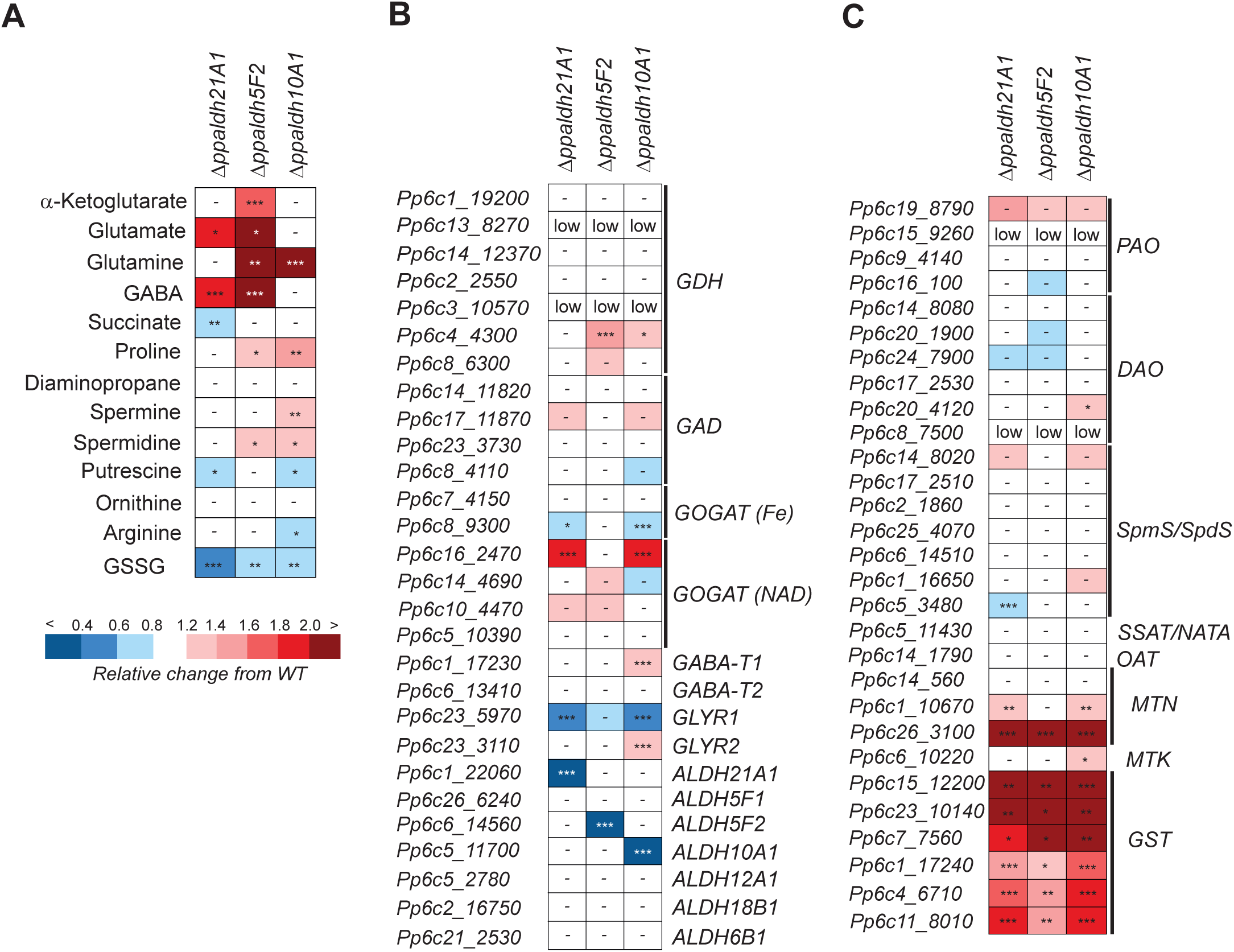
Metabolites and gene transcripts in three *P. patens aldh* knockouts versus WT. **(A)** Relative changes of selected metabolites compared to WT plants. Data were measured in triplicate. **(B, C)** Relative changes in levels of transcript expression determined by RNAseq analysis related to genes linked to GABA shunt and TCA cycle **(B)** and polyamine and GST metabolism **(C)**. Values are means of three biological replicates per genotype. Moss samples grown in the presence of 300 mM NaCl. Asterisks indicate statistically significant differences in knockout lines versus the controls in a paired Student’s t-test (t-test; *, **, and *** correspond to p-values of 0.05 > p > 0.01, 0.01 > p > 0.001, and p < 0.001, respectively).

Glu, GABA and succinate levels remained unchanged in the *aldh10A1* line (Fig. 6A). The *GABA-T1* gene was upregulated, which indicates faster conversion of GABA to SSAL, and both *glyoxylate/SSAL reductase (GLYR)* genes became dysregulated: *GLYR1* was downregulated and *GLYR2* was upregulated. Nevertheless, we observed a mild adjustment among polyamine-linked metabolites. Although no major changes in the expression of genes encoding Spd/Spm synthases (SpdS/SpmS), PAO and DAO were detected (Fig. 6C), levels of Arg and Put were slightly decreased, and those of Spd and Spm were slightly elevated.

Interestingly, while the *aldh21A1* knockout accumulated only more Glu but not glutamine (Gln), the *aldh10A1* line did the opposite. In contrast, the *aldh5F2* mutant accumulated both amino acids Glu and Gln. Although the expression of *GS* genes was not altered, *aldh21A1* and *aldh10A1* mutants showed differential expression of two *GOGAT* genes (Pp6c8_9300 and Pp6c16_2470) (Fig. 6B). Both *aldh5F2* and *aldh10A1* knockouts contained ∼ 40 - 50% more Pro, despite very similar expression of Pro biosynthetic genes. Finally, the expression of other *ALDHs* indirectly associated with the GABA shunt or Glu metabolism (e.g., *ALDH2A1*, *ALDH6B1*, *ALDH12A1*, and *ALDH18B1*) remained unchanged across all tested knockout lines.

## DISCUSSION

### Most of moss ALDH genes are less responsive to abiotic stresses compared to barley

The expression pattern of complete *ALDH* gene families in moss and barley under various abiotic stresses (Fig. 2) in this work is only in part well correlated with RNA-seq data available for *P. patens* at MAdLandExpression database (https://peatmoss.plantcode.cup.uni-freiburg.de/) and for *H. vulgare* at BarleyExpDB (http://barleyexp.com/), where similar conditions can found for drought (Hiss *et al*., 2014; Perroud *et al*., 2018; Cantalapiedra *et al*., 2017; Harb *et al*., 2020), salt, low or high temperatures (Beike *et al*., 2015; Perroud *et al*., 2018; Cantalapiedra *et al*., 2017; Schubert *et al*., 2019; Pacak *et al*., 2016). However, the expression of many *ALDH* genes is conflicting between different published datasets or is even missing. Moreover, available RNA-seq data do not show any significant responses to gibberellin or strigolactone treatments except for the *PpALDH21A* gene, which is upregulated ∼ 2-fold by GA9 methyl ester (Hiss *et al*., 2014), whereas it is downregulated 4-fold by strigolactone analog G24 (Perroud *et al*., 2018). Thus, these two hormones were not included in our experiment.

A direct correlation between the number of stress-responsive elements in promoters (Supplementary Fig. S2) and expression changes (Fig. 2) was not found. High correlation was observed between the number of stress-responsive elements and positive responses in only a few genes, such as *PpALDH3H1*, *PpALDH11A2*, *PpALDH21A1, HvALDH3E2* and *HvALDH5F1.* On the contrary, some genes with a high number of regulatory elements were weakly responsive to stress.

When comparing responses in moss and barley, a major difference was observed within the *ALDH7* family. Whereas the barley *ALDH7B6* is strongly induced by drought, salinity, MeJA, ABA, and cytokinin, the moss *ALDH7B4* gene is unresponsive or even slightly downregulated. The ALDH7, also known as Δ^1^-piperideine-6-carboxylate (P6C) dehydrogenase, catalyzes the conversion of AASAL to α-aminoadipate in the lysine degradation (saccharopine) pathway, with P6C representing the cyclic Schiff base of AASAL. In seed plants, *ALDH7* expression is well known to be induced by dehydration, high salinity, or ABA treatment (Guerrero *et al*., 1990; Stroeher *et al*., 1995) and its overexpression or mutation can confer tolerance or sensitivity to abiotic stresses (Rodrigues *et al*., 2006; Kotchoni *et al*., 2006; Chen *et al*., 2015; Shin *et al*., 2009). Since α-aminoadipate is further degraded to acetyl-CoA, which enters the TCA cycle or is used for lipid synthesis, *Physcomitrium patens* may not substantially benefit from lysine degradation during abiotic stress as higher plants do.

A similar difference between moss and barley appears in the *ALDH2* family. All five barley *ALDH2B* and *ALDH2C* genes are strongly responsive to drought, salinity, MeJA, ABA, cytokinin, or lack of nitrogen, whereas both moss *ALDH2B* genes remain unresponsive under the same conditions. The *ALDH2* genes from the ALDH2B subfamily were originally identified among genes that restored male fertility in maize (Liu *et al*., 2001, 2002). These enzymes produce acetate for acetyl-CoA biosynthesis. Thus, it seems that again, *Physcomitrium patens* does not produce acetyl-CoA in this ALDH-dependent pathway during abiotic stress. The ALDH2C subfamily, present in barley, comprises cytosolic members that oxidize longer aliphatic aldehydes, including *t*-2-hexenal and *t*-2-nonenal, from the lipoxygenase pathway, as well as aromatic aldehydes from the phenylpropanoid pathway (Končitíková *et al*., 2015). They modulate the levels of soluble and cell wall-linked ferulate/sinapate esters, affecting lignin biosynthesis (Nair *et al*., 2004; Mittasch *et al*., 2013; Yamamoto *et al*., 2024). Because true lignin has not been detected in bryophyta and appears only in vascular plants, *ALDH2C* genes are thus not needed and are absent.

Proline (Pro) was reported to accumulate early at the onset of salt stress in barley or wheat, while glycine betaine later on (Nakamura *et al*., 1996; Carillo *et al*., 2008). The upregulation of both *HvALDH18* genes, leading to Pro synthesis, is thus in good agreement with previous publications (Fig. 2). In contrast, *PpALDH18B1* was not stress-responsive in our experiments, although *Physcomitrium* accumulates Pro and soluble sugars in response to drought and salinity (Erxleben *et al*., 2012).

The higher number of *ALDH11* genes is typical of lower plants such as mosses or lycophytes, while seed plants carry only a single copy. The high variation in expression among the four *ALDH11* genes (Fig. 2) shows that *Physcomitrium* tries to cope with various stresses by fine-tuning glycolysis, the Calvin-Benson cycle and the pentose phosphate pathway. Non-phosphorylating GAPDH reaction catalyzed by ALDH11 (EC 1.2.1.9) produces NADPH (and no ATP) and bypasses two glycolytic reactions of NAD^+^-dependent GAPDH (EC 1.2.1.12) and phosphoglycerate kinase (EC 2.7.2.3), generating NADH and ATP. Moreover, neither *Physcomitrium* nor barley synthesizes reduced sugars, such as mannitol, as primary photosynthetic products, where ALDH11 is known to participate (Gao and Loescher, 2000). Barley *ALDH11* is downregulated in response to various stresses (Fig. 2), except during nitrogen starvation and cold stress. Downregulation mimics the Arabidopsis *aldh11* knockout (Rius *et al*., 2006), which is compensated by induced NAD^+^-dependent GAPDH and glucose-6P dehydrogenase to increase NADH and NADPH production, respectively. Secondly, *aldh11* knockout in Arabidopsis displayed reduced CO_2_ assimilation, and the gene coding for a cytosolic malate dehydrogenase, which regulates a redox balance, was strongly upregulated.

Among GABA shunt-associated *ALDHs*, transcription of *PpALDH21A1* increased upon drought, salt and ABA exposure. Previous studies performed in moss revealed that *PpALDH21A1* was upregulated upon osmotic and salt stresses and levels of Pro or sucrose were found elevated (Cuming *et al*., 2007; Wang *et al*., 2008). On the contrary, transcription of both *PpALDH5* is not drought-responsive; *PpALDH5F1* transcripts increased only upon salt stress. In barley, lacking the *ALDH21* gene, the stress-responsiveness was shifted towards the only *HvALDH5F1* gene, which is highly upregulated by drought, salt and ABA. It has been reported that GABA accumulates upon high nitrate in closely related wheat (Woodrow *et al*., 2017). However, in our work, neither of the two HvALDH10 genes responded to the excess of nitrogen, and thus, the GABA accumulation likely appears due to higher GAD activity.

### *ω*-Aminoaldehydes are oxidized by the ALDH10 whereas ALDH5 family prefers SSAL

In terms of substrate and coenzyme preferences, PpALDH10A1 resembles both the pea, maize and tomato ALDH10 family members (Tylichová et al., 2010; Kopečný et al., 2013), which are aminoaldehyde dehydrogenases (AMADHs). The active site of PpALDH10A1 contains Trp292 and Ile448, residues associated with the strong binding of APAL and ABAL, and consequently with AMADH activity. The poor BAL dehydrogenase (BADH) activity of PpALDH10A1 correlates well with the fact that *P. patens* does not accumulate glycine betaine (Erxleben *et al*., 2012).

Two barley AMADHs (HvALDH10A6 and HvALDH10A9), previously shown to oxidize APAL, ABAL, TMABAL, and BAL (Fujiwara *et al*., 2008), are very similar to maize AMADHs (Kopečný *et al*., 2013). HvALDH10A6 (Uniprot Q94IC0) is a peroxisomal enzyme, whereas HvALDH10A9 (Uniprot Q94IC1) resides in the cytosol (Nakamura *et al*., 2001). Their substrate channel comprises the aromatic residue Phe291/Trp289, which increases the preference for APAL or ABAL, as shown for maize isoforms. HvALDH10A6 contains Ile447 is thus AMADH (Fujiwara *et al*., 2008; Kopečný *et al*., 2013). In contrast, HvALDH10A9, which possesses a cysteine at the equivalent position, displays a BADH activity, which is in agreement with the fact that glycine betaine accumulates upon prolonged salinity stress in barley and wheat (Nakamura *et al*., 1996; Carillo *et al*., 2008).

Negatively charged and four carbon-containing SSAL was identified as the major substrate for both moss and barley ALDH5 members with *K*_m_ values of ∼ 70 - 90 μM (Table 3). Oxidation of other aldehydes was minimal. The barley enzyme has been briefly studied in the past and a *K*_m_ value of 15 μM for SSAL was reported (Yamaura *et al*., 1988). Its binding involves two residues, Arg169 and Arg287, together with Ser450 and Tyr115 (PpALDH5F1 numbering). In human ALDH5A1 (Kim *et al*., 2009), the SSAL was trapped in the active site *via* salt bridges with the equivalent Arg213 and Arg334, and *via* H-bond to Ser498. Although the active site of PpALDH21A1 (Kopečná *et al*., 2017) differs from that of ALDH5 (Fig. 4H), again, two Arg residues, Arg121 and Arg457, with Tyr296 bind the carboxylate group of SSAL. Positively charged ω-aminoaldehydes are thus repelled by the presence of two positively charged Arg side chains in the substrate channel of ALDH5/ALDH21, as evidenced by site-directed mutagenesis in this work (Table 3 and Fig. 4I). Several SSALDHs were previously studied from other species, including human, wheat, potato and Arabidopsis ALDH5 (Cash *et al*., 1978; Kang *et al*., 2005, Galleschi *et al*., 1978; Satyanarayan and Nair, 1989; Busch and Fromm, 1999) with *K*_m_ values between 7 and 16 μM for SSAL and an inhibition effect of SSAL upon higher concentrations. PpALDH21 displayed a *K*_m_ value of 181 ± 7 μM for SSAL with NADP^+^ (Kopečná *et al*., 2017).

### GABA and Glu accumulated in moss aldh5F2 *or* aldh21A1 *knockouts*

The GABA shunt and the TCA cycle can compensate for each other, as shown in GABA permease mutants (Michaeli *et al*., 2011). Here, the absence of two SSALDHs created a bottleneck in the *aldh5F2* or *aldh21A1* knockouts, resulting in the accumulation of GABA and its precursor Glu, with little or no impact on succinate (Fig. 6A). Elevated Glu can be interconverted to 2-ketoglutarate by GDH or transaminases to enter the TCA cycle. It can also be recycled in the Gln synthase/Gln-oxoglutarate aminotransferase (GS/GOGAT) cycle, a key process in nitrogen assimilation, or used in other pathways such as Pro synthesis. The *aldh5F2* knockout contained twice the amount of Gln and 50% more of 2-ketoglutarate. Notably, one of the *GDH* genes (Pp6c4_4300) was upregulated in this knockout. However, similar levels of succinate suggest that the remaining SSALDHs can compensate for the effect of a single *aldh5* or *aldh21* knockout. Accumulation of GABA is in line with previous studies on Arabidopsis *aldh5* knockout, where blocked SSAL oxidation resulted in the accumulation of GABA, as well as GHB, while succinate levels were reported to decrease (Fait *et al*., 2005; Bouché *et al*., 2003; Toyokura *et al*., 2011). A double mutant lacking both *ALDH5 (SSADH)* and *GABA-T* (*POP2*) accumulated even higher levels of GABA while GHB levels were reduced (Ludewig *et al*., 2008). Unfortunately, due to the limitations of the used method, GHB levels were not quantified in our moss samples; thus, the contribution of GLYR enzymes couldnot be evaluated.

### ALDH10A1 contributed only negligibly to the GABA shunt in moss

The moss *aldh10A1* knockout accumulated more polyamines Spd and Spm than WT, confirming that while polyamine degradation (oxidation) is slowed, GABA levels were fully compensated by Glu decarboxylation *via* GAD (Fig. 6A). This finding indicates a minor flux of GABA from polyamines mediated by ALDH10A1 in *Physcomitrium patens* grown in media with 300 mM NaCl. Unchanged levels of more distant metabolites, namely 2-oxoglutarate and succinate, indicated an unaffected TCA cycle. In contrast, Arabidopsis single knockouts *aldh10A8* and *aldh10A9* (Missihoun *et al*., 2011) as well as the *aldh10A8/aldh10A9* double mutant (Jacques *et al*., 2020), contained decreased GABA levels compared with WT under salt stress at 150 mM NaCl. Thus, both *ALDH10* genes contribute to the GABA-shunt pathway. In tea (*Camellia sinensis*) plants, *amadh1* (*aldh10*) and *cuao1* (*dao1*) antisense mutants contained 2-fold lower GABA levels than WT under drought stress (Cao *et al*., 2024). Thus, ALDH10 appears to contribute little or not at all to the GABA shunt in moss under salt stress (and others), in contrast to higher plants. This is consistent with the unchanged expression of *ppaldh10A1* under various stress conditions.

### Upregulated glutathione-S-transferase genes compensate for the missing ALDHs in moss

Our moss *aldh* mutants displayed slower growth than WT. Consistently, *ALDH7B2* overexpression improved, while *aldh7B4/aldh3I1* knockouts impaired photosynthetic performance under salt stress (Gautam et al., 2020; Missihoun et al., 2018). In our moss *aldh* knockouts, although the gene that encodes cytochrome *b_6_*was downregulated, the function of PSI was not altered and the photosynthetic efficiency of PSII was only negligibly affected. However, the *aldh5, aldh10* and *aldh21* mutants had a visibly reduced ability to induce NPQ, which is associated with a higher sensitivity to photo-oxidative damage under high light conditions. Indeed, light-induced cell death and accumulation of reactive oxygen species were reported in Arabidopsis *aldh5* knockout (Bouché *et al*., 2003).

ALDHs detoxify lipid peroxidation products such as malondialdehyde, crotonaldehyde, methylglyoxal, 4-hydroxy-2-hexenal (4-HHE), and 4-hydroxy-2-nonenal (4-HNE), as their increased concentration can damage proteins and DNA and lead to cell death, and levels of the above are decreased in *ALDH* overexpressors (Sunkar *et al*., 2003; Kotchoni *et al*., 2006). Reactive carbonyl compounds are also detoxified by GSTs that catalyze the conjugation of reduced glutathione (GSH) to various electrophiles, including 4-HHE or 4-HNE via Michael-type reaction. Consequently, the conjugate is exported from the cell, and thus, the total glutathione content (GSH + GSSG) can change significantly. The above aldehydes were reported to induce the expression of several GST genes in Arabidopsis (Alméras *et al*., 2003) as well as in pumpkin, leading to up to a 10-fold increase in GST activity (Fujita and Hossain, 2003).

Transcriptomic analysis of all three *aldh* knockout lines prepared in this work, namely *aldh5, aldh10* and *aldh21* mutants, revealed several upregulated *GST* genes (Fig. 6C). In *Physcomitrium*, there are at least 37 genes of a *GST* superfamily sorted into ten classes (Liu *et al*., 2013). The tree upregulated genes Pp6c15_12200, Pp6c23_10140 and Pp6c7_7560 in all three *aldh* knockouts belong to “phi” class encoding PpGSTF1, PpGSTF6 and PpGSTF7. The other upregulated genes, Pp6c1_17240 and Pp6c4_6710, belong to the “zeta” and “Ure2p“ classes, respectively, and encode PpGSTZ1 and PpUre2p1 enzymes. Phi- and zeta-GSTs contain a serine residue in their active site known to promote GSH-conjugation and/or peroxide reduction reactions and overexpression of several GSTs was shown to confer tolerance to drought and salt stresses (Sylvestre-Gonon *et al*., 2019).

The induction of multiple *GST* genes in all three *aldh* moss knockouts suggests higher GST activity and it may represent a compensatory protective mechanism to detoxify elevated reactive aldehydes when lacking functional ALDH. Increased conjugation is supported by the decreased GSSG levels in moss *aldh* knockouts because the Michael-type addition catalyzed by GSTs is linked to conjugation of GSH with aldehydes but not GSH oxidation to GSSG. Unfortunately, our method did not allow for the quantification of total glutathione; however, we suggest that the consumed GSH will be compensated for by the reduction of GSSG to GSH, thereby maintaining the glutathione redox state. Indeed, the total glutathione pool was found to be decreased in the double *aldh7B4/aldh3I1* knockout in Arabidopsis (Missihoun *et al*., 2018), but no transcriptomic data were analyzed in that study.

We can conclude that comparative analysis of aldehyde dehydrogenase families in moss and barley revealed divergent responses to abiotic stress and hormones. Moss mutants allowed us to find that cytosolic SSALDH (ALDH21A1), unique to mosses, contributes to GABA-shunt in a fashion similar to a mitochondrial SSALDH (ALDH5F2). On the contrary, ALDH10, known to contribute significantly to the GABA shunt in higher plants under salt stress, appears to contribute only negligibly to the GABA shunt in moss. Final transcriptomic analysis points to upregulation of multiple *GST* genes in moss *aldh* mutants and uncovers a novel metabolic crosstalk between ALDHs and GSTs.

**ABBREVIATIONS**
AASAL: α-aminoadipate-semialdehyde;
ABA: abscisic acid;
ABAL: 4-aminobutyraldehyde;
ALDH: aldehyde dehydrogenase;
APAL: 3-aminopropionaldehyde;
AMADH: aminoaldehyde dehydrogenase;
BA: benzyl adenine;
β-ALA: β-alanine;
DAO: diamine oxidases;
GABA: γ-aminobutyric acid;
GABA-T: transaminase;
GAD: glutamate decarboxylase;
GBAL: 4-guanidinobutyraldehyde;
GHB: 4-hydroxybutyrate;
GDH: glutamate dehydrogenase;
GRSAL: glutaric semialdehyde;
GSAL: glutamate-γ-semialdehyde;
Hv: *Hordeum vulgare*;
MeJA: methyl jasmonate;
MST: microscale thermophoresis;
NiR: nicotinamide riboside;
PAO: polyamine oxidase;
PEG: polyethylene glycol;
Pp: *Physcomitrium patens*;
SSAL: succinic semialdehyde;
TCA: tricaboxylic acid;
TMABA: trimethyl-γ-aminobutyric acid;
TMABAL: trimethyl-4-aminobutyraldehyde

## SUPPLEMENTARY DATA

**Supplementary Table S1.** Primers and dual-labeled probes used in qPCR analyses of moss *ALDH* genes.

**Supplementary Table S2.** Primers and dual-labeled probes used in qPCR analyses of barley *ALDH* genes.

**Supplementary Table S3.** Primer pairs used for ALDH cloning.

**Supplementary Table S4.** Data collection and refinement statistics.

**Supplementary Table S5.** Primers used to clone 5’- and 3’-fragments into pBNRF vector and primer pairs used for genotyping of transformants.

**Supplementary Table S6.** Detailed gene nomenclature of *Physcomitrium patens ALDH* superfamily.

**Supplementary Table S7.** Detailed gene nomenclature of *Hordeum vulgare ALDH* superfamily.

**Supplementary Table S8.** Specific activities of ALDH5 variants with SSAL.

**Supplementary Table S9.** Levels of metabolites in moss WT and three *aldh* knockouts.

**Supplementary Fig. S1.** Gene annotation corrections and relative expression levels of *ALDH* superfamily genes in *Physcomitrium patens* and *Hordeum vulgare*.

**Supplementary Fig. S2.** Promoter analysis of cis-regulatory elements.

**Supplementary Fig. S3.** Gel permeation chromatography of moss ΔPpALDH10A1 and PpALDH5F1.

**Supplementary Fig. S4.** Sequence alignment of ALDH5 enzymes analyzed in this work with human ALDH5A1.

**Supplementary Fig. S5.** Example of phenotyping and genotyping of *Physcomitrium aldh5F2* knockout mutants.

**Supplementary Fig. S6.** Volcano plots illustrating differentially expressed genes in three moss *aldh* mutants vs WT in RNA-seq data.

**Supplementary Fig. S7.** A heat map of DEGs in moss *aldh5F2* mutant vs control generated from RNA seq data reflecting log(e)-transformed transcripts per million (TPM) values.

**Supplementary Fig. S8.** A heat map of DEGs in moss *aldh21A1* vs control generated from RNA seq data reflecting log(e)-transformed transcripts per million (TPM) values.

**Supplementary Fig. S9.** A heat map of DEGs in moss *aldh10A1* vs control generated from RNA seq data reflecting log(e)-transformed transcripts per million (TPM) values.

**Supplementary Fig. S10.** Photosynthetic control parameters in moss *aldh* mutants during light exposure.

**Supplementary Dataset S1.** Assembled ALDH sequences in moss and barley.

**Supplementary Dataset S2.** DEGs identified in moss *aldh* mutants.

## ACKNOWLEDGMENTS

We acknowledge SOLEIL for the provision of synchrotron radiation facilities (proposal IDs 20191181 and 20210831) in using PROXIMA beamlines and we thank the staff for assistance in using the beamlines.

## AUTHOR CONTRIBUTIONS

DK, MK, VB and SM designed the research; DJK and JB performed a real-time qPCR and the phenotyping analysis; MK, JB, RaK: prepared enzyme variants and their mutants, and measured enzyme properties with kinetics; DJK and KvS prepared moss knockout lines; DJK, DK, JB, MK, MV and KM performed and evaluated RNA-seq analyses; DJK and RoK analyzed photosynthetic parameters, KK and SCZ were involved in metabolite measurements; AV, DK and SM performed the crystallographic study and analyzed the crystal structures; DK and SM wrote the paper. All authors reviewed the results and approved the final version of the manuscript submitted for publication.

## DATA AVAILABILITY

The atomic coordinates and structure factors have been deposited in the Protein Data Bank (www.wwpdb.org) under accession codes 8OF3 for the apoform of PpALDH5F1, 8OFM for the PpALDH5F1 complex with NAD^+^ in an extended conformation and 8OF1 for the PpALDH5F1 complex with NAD^+^ in a contracted conformation. RNA-seq RAW datasets of moss aldh knockouts and WT were deposited in the NCBI Gene Expression Omnibus (GEO) database (https://www.ncbi.nlm.nih.gov/geo/) under accession code GSE299690. Differentially expressed genes from RNA-seq, qPCR data, metabolite, kinetic and ligand binding data were deposited in the Zenodo repository at https://doi.org/10.5281/zenodo.18186873. Other data are available in the Supplementary data.

