## Supplemental Table S1 for "Insight into GABA shunt-associated aldehyde dehydrogenases (ALDH) and stress responses of ALDH superfamily in moss and barley"

**Supplementary Table S1. Primers and dual-labeled probes used in qPCR analyses of moss *ALDH* genes.**

| Target gene | Sequences of the dual-labeled probe and primers |
|---|---|
| ***PpEFα*** | [6FAM]CCTGCGGTTGCCCCTTCAGGA[TAM], CAAGAGGCCATCCGACAAAC, CAGTTCCAATACCTCCAATCTTGTAG |
| ***PpUBQ2*** | [6FAM]TGCCTTCAGGACCAAGGTTTTCCACC[TAM], CCATTCAAGCCCCCTAAGG, AAATGCTCCCATTGCTATTAATGTT |
| ***PpALDH2B1*** | [6FAM]CGGCAGCTCAGAGCAACCTTAAACCA[TAM], AGTGGGACGTCAAATTATGCAA, TTGCCTCCAAGTTCAAGATTCA |
| ***PpALDH2B2*** | [6FAM]CCCTTTCCAGCGCCAGCGAA[TAM], ACCCTGGCCGAAAATGC, CGAGCAAATCTGCGTACTTGAG |
| ***PpALDH3H1*** | [6FAM]CCGACCTCGGCAAGCCCTCG[TAM], CGAAATCGTGCAGACCCTGTA, AGGGACACCTCGGTGACGTA |
| ***PpALDH3K1*** | [6FAM]AGCTGCTGCAAAGCACCTCACGC[TAM], GCCAAGGTGGGCAGGATTA, AAGGGACACTTTCCTCCAAGCT |
| ***PpALDH3K3*** | [6FAM]AGCAATCTCAGCCGGTTGCGCT[TAM], TGGCGGTAGATCCCCTGAT, CGACGCCTTCAGGCAAA |
| ***PpALDH3K2*** | [6FAM]CCTTTCCTCTTGGCGGTAGATCCCG[TAM], TCGTCATCGCTCCATGGAA, CAACCGGCTGCAATTGCT |
| ***PpALDH3H2*** | [6FAM]CACGCCGGCTCTACTGGCCAA[TAM], GCCTTCGGAAGTCGTTTACG, AGTTGTCCATGTAGAGAGGGATGAG |
| ***PpALDH5F2*** | [6FAM]CACCTCTTATGCGCTTCCAGACGGA[TAM], AGAGAGGAGGTTTTTGGTCCTGTA, ATTTGCCATCTTGATAGCCTCTTC |
| ***PpALDH5F1*** | [6FAM]TATTTGGGCCAGTTGCACCCCTTGT[TAM], ACGCTAGTGACGAAATGCTTATCTT, TGCTTCCTCATCTGTGTTAAATCG |
| ***PpALDH6B1*** | [6FAM]CATGCCAGACGCAGATCCCGAA[TAM], TGCGAAAAACCATGCTGTGA, TCCGACAAGAGCATTCAATGTT |
| ***PpALDH7B4*** | [6FAM]TACCACCTGCCGGCGCCTG[TAM], TGGTACTGCCGGTCAACGTT, CATCATAGACTTTCTCGTGCACAA |
| ***PpALDH10A1*** | [6FAM]ATGCCCCTCTCGAACTCCCCATG[TAM], GCGGAGAAATTAGACGAGAGACA, GCAAGATGTTGCACTTGAACTGT |
| ***PpALDH11A1*** | [6FAM]CCGTATCGAAAATGGGTCCAGCCC[TAM], CACCTTTCAATTATCCTGTCAACCT, CAACTGCATTCCCAGCAATG |
| ***PpALDH11A2*** | [6FAM]CAAGGATCAGAAAAACGCCATCGCG[TAM], TTCACAAGTTCGCTGGGATTC, TTGCAATCTCCTTCACAAGCA |
| ***PpALDH11A4*** | [6FAM]TCCTTGCCATTCCCCCATTTAACTACCC[TAM], TGCCTTGCTTCGAAGATTCC, GGACACGGCCAGGTTGAC |
| ***PpALDH11A3*** | [6FAM]ACTGCCTTGCATCGAAGATTCCTCTTGGT[TAM], CCCAGGAAACGGAAGAAATAAA, GTAGTTGAATGGCGGAATAGCAA |
| ***PpALDH12A1*** | [6FAM]ACTCATCCAGCGGGTTGCACCAA[TAM], AACCTCAGGTCGAGACGTTCTT, ACCGCCTGGGCAAAGC |
| ***PpALDH18B1*** | [6FAM]TTGGCCGTCTCACTATCAAACCTGGG[TAM], GGCGTGTCGAAGGCTTTG, TCCGGAGAGAACTGGCAAGT |
| ***PpALDH21A1*** | [6FAM]CCGCATTTATGGCGAGCACATTCC[TAM], CAGGTTGCAGCTGAAGAAAGTG, TGTTCCGTGCAGAGATATCCAA |
| ***PpALDH23A1*** | [6FAM]CGAAGCAAGCTGCTCCAAATCTTGTCA[TAM], TCACGGGCTCGAACAAGAC, TTTCCTCCGAGCTCCAATTG |
