## Supplemental Table S2 for "Insight into GABA shunt-associated aldehyde dehydrogenases (ALDH) and stress responses of ALDH superfamily in moss and barley"

**Supplementary Table S2. Primers and dual-labeled probes used in qPCR analyses of barley *ALDH* genes.**

| Target gene | Sequences of the dual-labeled probe and primers |
|---|---|
| ***HvEF2*** | [6FAM]AGCAAGTCCCCCAACAAGCATAACCG[TAM], AAGTCCTGCCGCACTGTCAT, GGGCGAGCTTCCATGTAAAG |
| ***HvACT*** | [6FAM]AAGAGTTCCCTCCACCTCCCCGACC[TAM], ATCCTCGGCTTCTCATTCCA, CCAAGTATCCTCCAGCATGTCA |
| ***HvALDH2B1*** | [6FAM]ACAGCTCCTCATCAACGGCAAATTCG[TAM], ATCTCGCCTTCTGTCCAAGTC, AAGTCTTGCCAGATGCAGCAT |
| ***HvALDH2B5*** | [6FAM]TCCAAGGCTCGCGCCCAGC[TAM], TGCACGAGAGCGTGTACGA, TGAACTGCTGGCCATCGAT |
| ***HvALDH2C1*** | [6FAM]AGCCCGCCATCTTCACCGACG[TAM], GCAGCGACAAGGGTTACTACATC, CGCAATCGACATGTCATCCT |
| ***HvALDH2C2*** | [6FAM]AGATCCGTTCGATGTGGCCACCA[TAM], GCTGCCCAGAACTGGAAAGTC, CTCAAATTGCTCCTTGTCAACCT |
| ***HvALDH2C3*** | [6FAM]AACTCGCCATCAGCGCCAACTTCTT[TAM], CGACGCGGACGTAGACATAG, CAATGCAAGCTTCTCCCTTGT |
| ***HvALDH3E1*** | [6FAM]ACCCGCCGTGGAACGACGAA[TAM], GCTTCCTGGTGGAGTTCATGTT, CGCAGCAACCCGAGTTTG |
| ***HvALDH3E3*** | [6FAM]ACTCTCTGGCGCACTAGCAGCTGGC[TAM], GCCACTAGGTTTGGCTTTGG, CGGAGGGCTTTACAACTACAACA |
| ***HvALDH3E2*** | [6FAM]TGGCGGCATCCATCCTGCAC[TAM], CATTTCCAGCGGCTGAGTAAC, ACAGGTTCTTGGCGTCTAAGTTG |
| ***HvALDH3H1*** | [6FAM]TATCGGCAGACTTGTCCCGCGTC[TAM], CAAGTTCTACGGAAAGGATCCACTT, CCTGTTGAAATGGTCGACATTC |
| ***HvALDH3H2*** | [6FAM]TCCTTTCTTGCTATCTATCGAGCCCGTCA[TAM], TCTCGTCATATCAGCCTGGAACT, CAGCATTCCCAGCAGCAATT |
| ***HvALDH5F1*** | [6FAM]TCCCCCAACTCTAGCTGATCGCAGACT[TAM], GCAAAGCGTGTGTATGGTGATATTA, TACCCCAATAGGCTGCTTCAG |
| ***HvALDH6B1*** | [6FAM]CATTGAATGCCCTTATTGCTGCCGG[TAM], CCTTCCTGATGCTGATCGAGAT, GCCCGGCTGCACCAA |
| ***HvALDH7B6*** | [6FAM]CATCATACCATCTGAACGCCCAAACCA[TAM], GGCTAAGTCGTCAGCTAAATGGAT, AGAGGATTCCAGACCTCCATCA |
| ***HvALDH10A9*** | [6FAM]TCATCACACCCTGACGTCGACAAGG[TAM], GGTAATGAAGCTGGTGCTCCTTT, GCATAGCTCCCGGTAAATGC |
| ***HvALDH10A6*** | [6FAM]ATGCACCAATCTCTCTGCCCATGGA[TAM], GCTGAAGCCTTAGATGGAAAACA, ACCCCGATGGGTTCTTTAAGA |
| ***HvALDH11A3*** | [6FAM]CGCGGTGGCAGCCCTCCA[TAM], TTGTGCTCAAGCCTCCAACTC, ACCGGCCAGGTGAAAGC |
| ***HvALDH12A1*** | [6FAM]TGCTCTGCCCAGTCTATGCTATTCATGCA[TAM], TTACGCTTGCAGTGGTCAGAA, CAAGTAGCCCACTAGACGACCAA |
| ***HvALDH18B1*** | [6FAM]TTTTTGAGTCCCGACCTGATGCCTTG[TAM], GCCCTTTGGGTGTTCTATTGAT, CGAATGGCTAAAGACGCAATC |
| ***HvALDH18B2*** | [6FAM]CACCGATCGCGATTTCCGGG[TAM], TGTGACATCGTCTCAGCTTCTTG, CTGGTGCCCAAAACTTGGA |
| ***HvALDH22A1*** | [6FAM]ATGATCCCCAAGCCTATTCAGTATCCCGTC[TAM], GTGGTGGCCATACGTCAAAAC, TGCTGGAATGCAAACCCATT |
