## Supplemental Table S3 for "Insight into GABA shunt-associated aldehyde dehydrogenases (ALDH) and stress responses of ALDH superfamily in moss and barley"

**Supplementary Table S3. Primer pairs used for selected *ALDH* cloning.**

| Primer name | Primer sequence |
|---|---|
| *PpALDH5F2* | 5´-CACTCATATGGGGTTTGCGCAAGTTGGT-3´ |
| | 5´-CACGTCGACTCAGCACATGAAGGGTTGCT-3´ |
| *PpALDH5F1* | 5´-CACTCATATGGGCACAATGGTGCAGC-3´ |
| | 5´-CATGTCGACTCAACCGACGGGTTGTG-3´ |
| *HvALDH5F1* | 5´-ACAGGATCCGATGGGCAGCGTGGACGC-3´ |
| | 5´-TGTCTCGAGTCAACCCAGGTTGCCCATGC-3´ |
| *PpALDH10* | 5´-CATGGATCCGATGGGTCTTCACGCCG-3´ |
| | 5´-CACCTCGAGTTACAACTTGGAAGGCTTGG-3´ |
| *Y115A – PpALDH5F1* | 5´-GCCGGTGCAGCCTTTGTAGAGTATT-3´ |
| | 5´-TACGATCTCTCCGACTGCTTCA-3´ |
| *R169A – PpALDH5F1* | 5´-P-GCCAAGGTAGCTCCAGCGTTAG-3´ |
| | 5´-P-AGTAATCATGGCCAGAGGAAAAT-3´ |
| *R287A – PpALDH5F1* | 5´-P-GCGAACTCTGGCCAAACTTGTG-3´ |
| | 5´-P-GTACTTGCCGGCCAACACTCCTT-3´ |
| *S450A – PpALDH5F1* | 5´-P-GCAACTGAGGTGGCACCTTTCG-3´ |
| | 5´-P-AATGAGGCCTTCATTTAGG-3´ |
