## Supplemental Table S4 for "Insight into GABA shunt-associated aldehyde dehydrogenases (ALDH) and stress responses of ALDH superfamily in moss and barley"

**Supplementary Table S4. Data collection and refinement statistics.**

| | **PpALDH5F1 apoform** | **PpALDH5F1 with contracted NAD$^+$** | **PpALDH5F1 with extended NAD$^+$** |
|---|---|---|---|
| PDB code | 8OF3 | 8OF1 | 8OFM |
| Ligand | - | NAD$^+$, sulfate | NAD$^+$, sulfate |
| Cryoprotection | PEG 400 | PEG 400 | Glycerol |
| Space group | H32 | H32 | H32 |
| Asymmetric unit | 2 monomers | 2 monomers | 2 monomers |
| Unit cell (Å) | | | |
| a | 212.7 | 212.50 | 212.31 |
| b | 212.7 | 212.50 | 212.31 |
| c | 186.7 | 186.74 | 187.35 |
| α (°) | 90.0 | 90.0 | 90.0 |
| β (°) | 90.0 | 90.0 | 90.0 |
| γ (°) | 120.0 | 120.0 | 120.0 |
| Resolution limits (Å)[a] | 1.62, 1.62, 1.3 | 2.5, 2.5, 1.8 | 2.1, 2.1, 1.8 |
| Resolution (Å) | 131.1 – 1.34 (1.5 – 1.34) | 131.1 – 1.81 (2.0 – 1.81) | 131.2 – 1.83 (2.0 – 1.82) |
| Observed reflections | 4989281 (195992) | 1612046 (61149) | 2154693 (95828) |
| Unique reflections | 241670 (12083) | 77952 (3898) | 103740 (5187) |
| Completeness (%) | 68 (12.7) | 53.4 (10.1) | 73.4 (14.1) |
| I/σ (I) | 15.6 (1.8) | 13.9 (2.2) | 14.5 (1.7) |
| $R_{sym}$ (%) | 11.4 (226.6) | 13.2 (156.1) | 19.5 (220.7) |
| $R_{pim}$ (%) | 2.5 (56.3) | 2.9 (39.8) | 4.3 (51.6) |
| $CC_{1/2}$ | 0.998 (0.628) | 0.998 (0.838) | 0.998 (0.601) |
| $R_{cryst}$ (%)[b] | 14.9 | 17.2 | 17.3 |
| $R_{free}$ (%) | 16.4 | 19.4 | 19.6 |
| RMSD bond lengths (Å) | 0.012 | 0.009 | 0.008 |
| RMSD bond angles (°) | 1.18 | 1.07 | 1.08 |
| Average B (Å$^2$) | | | |
| Protein chain A/B | 20/19.6 | 35.9/31.5 | 31.3/32 |
| Water molecules | 35.1 | 41.1 | 43.9 |
| NAD$^+$ chain A/B | - | 53.9/43.2 | 55.7/52.7 |
| Ramachandran statistics (%)[b] | | | |
| Favored | 97.6 | 98.1 | 97.8 |
| Outliers | 0.0 | 0.0 | 0.0 |
| Clash score (PR)[b] | 1.32 | 1.90 | 2.2 |
| MolProbity score (PR)[b] | 0.98 | 1.21 | 1.04 |

Values for the highest resolution shell are in parentheses. $CC_{1/2}$= percentage of correlation between intensities from a random half-dataset.

[a]STARANISO (https://staraniso.globalphasing.org/cgi-bin/staraniso.cgi) applies an ellipsoidal mask: estimated resolution limits along the three crystallographic directions a*, b*, c*.

[b]Calculated with MolProbity (http://molprobity.biochem.duke.edu/)
