## Supplemental Table S5 for "Insight into GABA shunt-associated aldehyde dehydrogenases (ALDH) and stress responses of ALDH superfamily in moss and barley"

**Supplementary Table S5. Primers used to clone 5´- and 3´-fragments into the pBNRF vector and primer pairs used for genotyping of moss transformants**.

| Primer name | Primer sequence |
|---|---|
| | ***aldh5F2_casette1 (1120 bp)*** |
| *PpALDH5F2_SpeI_F1* | 5´-AAGAACTAGTAGAGAGAGAGAGCGAGGTCG-3´ |
| *PpALDH5F2_AgeI_R1* | 5´-ATGTACCGGTGTCTAAAAACGCCTAGTTAG-3´ |
| | ***aldh5F2_casette2 (1026 bp)*** |
| *PpALDH5F2_XhoI_F2* | 5´-GCGTCTCGAGCGATTTATTTCTTTGGTG-3´ |
| *PpALDH5F2_XbaI_R2* | 5´-AATTTCTAGAGTGAGTCGATCCCAAGTAGG-3´ |
| *Genotyping_casette1_5F2* | 5´-GTCCCCTCTACCACCTTCAC-3´, |
| | 5´-TGTTCGCCAGTCTTTACGG-3´ |
| *Genotyping_casette2_5F2* | 5´-GGTTCTTATAGGGTTTCGCTCAT-3´, |
| | 5´-ACCTTCGGCACAATTCAG-3´ |
| | ***aldh10A1_casette1 (1027 bp)*** |
| *PpALDH10A1_BamHI_F1* | 5´-CGCGGATCCCTTTCCAGTTGCTCGCATCC-3´ |
| *PpALDH10A1_XhoI_R1* | 5´-AAAACTCGAGATATCCCATTCAGCTTCATCGATGG-3´ |
| | ***aldh10A1_casette2 (1035 bp)*** |
| *PpALDH10A1_SpeI_F2* | 5´-GGAATACTAGTCTGTTGAATGGGCTATGTTTGGAGC-3´ |
| *PpALDH10A1_AscI_R2* | 5´-GGTGGGCGCGCCCAATGCGAGAGCTTCCTCTTCC-3´ |
| *Genotyping_casette1_10A1* | 5´-CTCGCTTCAAGGAGTGATGG-3´, |
| | 5´-GAGCTCGGTACCATAACT-3´ |
| *Genotyping_casette2_10A1* | 5´-GAGTTCTGTTAGGTCCTC-3´, |
| | 5´-CAGATAGACAATTGCAACAAGTCC-3´ |
| | ***aldh21A1_casette1 (1113 bp)*** |
| *PpALDH21A1_AvrII_F1* | 5´-TATCCCTAGGCTCTCAGTGTGAGCCAGTTCC-3´ |
| *PpALDH21A1_XhoI_R1* | 5´-TTATCTCGAGCGTCGATTGCATCATCTATGTCC-3´ |
| | ***aldh21A1_casette2 (1122 bp)*** |
| *PpALDH21A1_NotI_F2* | 5´-AATTCGCGGCCGCGTAACGCGCCATGCATTGTAG-3´ |
| *PpALDH21A1_SpeI_R2* | 5´-GCGCCACTAGTCTGGATACCGGAGTCTTTCAGG-3´ |
| *Genotyping_casette1_21A1* | 5´-AATTGGCTGCGCATGACG-3´, |
| | 5´-GGTTCTTATAGGGTTTCGCTCAT-3´ |
| *Genotyping_casette2_21A1* | 5´-GTATGAACTGTTCGCCAGTCTT-3´, |
| | 5´-AACATTCCGCATGACAAGAACC-3´ |
