## Supplemental Table S6 for "Insight into GABA shunt-associated aldehyde dehydrogenases (ALDH) and stress responses of ALDH superfamily in moss and barley"

**Supplementary Table S6. Detailed gene nomenclature of *Physcomitrium patens ALDH* superfamily.** The genome of *Physcomitrium patens* ecotype Gransden 2004 v 6.1 (Bi *et al*., 2024) was used for the *ALDH* sequence identification. Gene nomenclature follows that of the published closest monocot rice (Brocker *et al.*, 2013). *Corrected sequence based on sequence alignments comprising the same family from other plants. **Presence of stop codon in exon 9.

| **ALDH family** | ***ALDH* gene name** (Wood and Duff, 2009) | ***ALDH* nomenclature** (Brocker *et al.*, 2013) | **Phytozome ID** | **Exon #** | **AA #** |
|---|---|---|---|---|---|
| Family 2 | *PpALDH2A* | *PpALDH2B1* | Pp6c5_10180V6.1 | 11 | 553 |
| | *PpALDH2B* | *PpALDH2B2* | Pp6c4_16740V6.1 | 4 | 535 |
| Family 3 | *PpALDH3A* | *PpALDH3H1* | Pp6c22_7980V6.2 | 10 | 492* |
| | *PpALDH3B* | *PpALDH3K1* | Pp6c17_1850V6.1 | 10 | 479 |
| | *PpALDH3C* | *PpALDH3K3* | Pp6c15_1350V6.1 Pseudogene | 8 | 447 |
| | *PpALDH3D* | *PpALDH3K2* | Pp6c9_1290V6.1 | 8 | 485 |
| | *PpALDH3E* | *PpALDH3H2* | Pp6c19_6490V6.1 | 10 | 583 |
| Family 5 | *PpALDH5A* | *PpALDH5F2* | Pp6c6_14560V6.2 | 20 | 498 |
| | *PpALDH5B* | *PpALDH5F1* | Pp6c26_6240V6.2 | 19 | 492 |
| Family 6 | *PpALDH6A* | *PpALDH6B1* | Pp6c21_2530V6.1 | 4 | 574 |
| Family 7 | *PpALDH7A* | *PpALDH7B4* | Pp6c2_4640V6.2 | 15 | 511 |
| Family 10 | *PpALDH10A* | *PpALDH10A1* | Pp6c5_11700V6.1 | 15 | 559 |
| Family 11 | *PpALDH11A* | *PpALDH11A1* | Pp6c24_3410V6.1* | 10 | 497* |
| | *PpALDH11B* | *PpALDH11A2* | Pp6c24_5150V6.1 | 8 | 496 |
| | *PpALDH11C* | *PpALDH11A4* | Pp6c20_8570V6.1 | 8 | 496 |
| | *PpALDH11D* | *PpALDH11A3* | Pp6c23_3270V6.1 | 8 | 496 |
| | *PpALDH11E* | *PpALDH11A5* | Pp6c17_180V6.1** Pseudogene | 10 | 501** |
| Family 12 | *PpALDH12A* | *PpALDH12A1* | Pp6c5_2780V6.2 | 16 | 571 |
| Family 18 | *PpALDH18A* | *PpALDH18B1* | Pp6c2_16750V6.2 | 20 | 745 |
| Family 21 | *PpALDH21A* | *PpALDH21A1* | Pp6c1_22060V6.2 | 8 | 497 |
| Family 23 | *PpALDH23A* | *PpALDH23A1* | Pp6c24_1400V6.1 | 2 | 494 |
