## Supplemental Table S7 for "Insight into GABA shunt-associated aldehyde dehydrogenases (ALDH) and stress responses of ALDH superfamily in moss and barley"

**Supplementary Table S7. Detailed gene nomenclature of *Hordeum vulgare ALDH* superfamily**. The genome of barley cv. MorexV3 (Mascher *et al.*, 2021) was used for the *ALDH* sequence searches. Gene nomenclature follows that of the published closest monocot rice (Brocker *et al.*, 2013). *Corrected sequence based on sequence alignments comprising the same family from other plants.

| Barley ALDH family | Gene name | Gene *ALDH* nomenclature | Phytozome ID | Exon # | AA # |
|---|---|---|---|---|---|
| Family 2 | *HvALDH2A* | *HvALDH2B1* | HORVU.MOREX.r3.7HG0671360.1 | 11 | 549 |
| | *HvALDH2B.1* | *HvALDH2B5* | HORVU.MOREX.r3.6HG0612050.1 | 8* | 543* |
| | *HvALDH2B.2* | *HvALDH2B6* | HORVU.MOREX.r3.6HG0611990.1* | 8* | 543* |
| | *HvALDH2B.3* | *HvALDH2B7* | HORVU.MOREX.r3.6HG0611990.1* | 8* | 543* |
| | *HvALDH2C* | *HvALDH2C1* | HORVU.MOREX.r3.3HG0308840.1 | 7 | 500 |
| | *HvALDH2D* | *HvALDH2C2* | HORVU.MOREX.r3.1HG0069410.1 | 9 | 499 |
| | *HvALDH2E* | *HvALDH2C3* | HORVU.MOREX.r3.7HG0714980.1 | 8 | 513 |
| Family 3 | *HvALDH3A* | *HvALDH3E1* | HORVU.MOREX.r3.6HG0599280.1 | 9 | 485 |
| | *HvALDH3B* | *HvALDH3E3* | HORVU.MOREX.r3.6HG0599310.1 | 10 | 484 |
| | *HvALDH3C* | *HvALDH3E2* | HORVU.MOREX.r3.2HG0185000.1 | 9 | 492 |
| | *HvALDH3D* | *HvALDH3H1* | HORVU.MOREX.r3.5HG0455570.1 | 10 | 480 |
| | *HvALDH3E* | *HvALDH3H2* | HORVU.MOREX.r3.4HG0347330.1 | 10 | 489 |
| Family 5 | *HvALDH5* | *HvALDH5F1* | HORVU.MOREX.r3.6HG0570900.2 | 19 | 526 |
| Family 6 | *HvALDH6* | *HvALDH6B1* | HORVU.MOREX.r3.2HG0198720.1 | 19 | 536 |
| Family 7 | *HvALDH7* | *HvALDH7B6* | HORVU.MOREX.r3.5HG0480430.1 | 14 | 509 |
| Family 10 | *HvALDH10A* | *HvALDH10A9* | HORVU.MOREX.r3.6HG0626440.1 | 15 | 549 |
| | *HvALDH10B* | *HvALDH10A6* | HORVU.MOREX.r3.2HG0173750.1 | 15 | 506 |
| Family 11 | *HvALDH11* | *HvALDH11A3* | HORVU.MOREX.r3.2HG0130620.1 | 9 | 496 |
| Family 12 | *HvALDH12* | *HvALDH12A1* | HORVU.MOREX.r3.1HG0081000.1 | 15 | 551 |
| Family 18 | *HvALDH18A* | *HvALDH18B1* | HORVU.MOREX.r3.1HG0073870.1 | 19 | 716 |
| | *HvALDH18B* | *HvALDH18B2* | HORVU.MOREX.r3.3HG0302060.1 | 20 | 730 |
| Family 22 | *HvALDH22* | *HvALDH22A1* | HORVU.MOREX.r3.2HG0110060.1 | 14 | 594 |
