## Supplemental Table S8 for "Insight into GABA shunt-associated aldehyde dehydrogenases (ALDH) and stress responses of ALDH superfamily in moss and barley"

**Supplementary Table S8. Specific activities of ALDH5 variants with SSAL.** Measured in 150 mM HEPES buffer, pH 8.0, using 50 µM SSAL and 1.5 mM $NAD^+$ by monitoring the NADH formation at 30 °C. Error values stand for S.D. (n=4).

| **Enzyme** | **Specific activity** (nmol $s^{-1}$ $mg^{-1}$) | **Relative activity** (%) |
|---|---|---|
| HvALDH5F1 | 140.6 ± 8.0 | - |
| PpALDH5F2 | 15.5 ± 1.0 | - |
| PpALDH5F1-WT | 48.2 ± 1.7 | 100.0 |
| PpALDH5F1-Y115A | 2.7 ± 0.1 | 5.6 |
| PpALDH5F1-R169A | 3.9 ± 0.1 | 8.0 |
| PpALDH5F1-R287A | 1.1 ± 0.1 | 2.2 |
| PpALDH5F1-S450A | 15.5 ± 0.5 | 32.2 |
