## Supplemental Table S9 for "Insight into GABA shunt-associated aldehyde dehydrogenases (ALDH) and stress responses of ALDH superfamily in moss and barley"

**Supplementary Table S9. Levels of metabolites in moss WT and three *aldh* knockouts.** Moss samples were grown in the presence of 300 mM NaCl. Polyamines/diamines were purified on the Acquity UPLC BEH AMIDE column, whereas tricarboxylic, dicarboxylic and amino acids were purified on the ARION Polar C18 HPLC column. All values are given in nmol $g^{-1}$ DW as a mean ± S.D. (n=3). Asterisks indicate statistically significant differences in knockout lines versus the controls (non-treated plants) in a paired Student's t-test (t-test; *, **, and *** correspond to P-values of 0.05 > p > 0.01, 0.01 > p > 0.001, and p < 0.001, respectively).

| **Metabolite (nmol $g^{-1}$ DW)** | **Wild type** | ***Δppaldh5F2*** | ***Δppaldh10A1*** | ***Δppaldh21A1*** |
|---|---|---|---|---|
| a-Ketoglutarate | 124.4 ± 15.2 | 201.1 ± 24.2* | 131.4 ± 19.3 | 122.8 ± 29.0 |
| Glutamate | 807.6 ± 184.4 | 2109.6 ± 170.4* | 929.1 ± 197.3 | 1489.3 ± 182.1* |
| Glutamine | 316.6 ± 82.8 | 655.5 ± 140.2** | 842.7 ± 126.2*** | 321.2 ± 91.7 |
| GABA | 143.6 ± 25.8 | 321.4 ± 40.4** | 147.3 ± 19.4 | 281.1 ± 58.0*** |
| Succinate | 231.3 ± 27.5 | 236.2 ± 66.7 | 216.5 ± 51.6 | 183.6 ± 19.5** |
| Proline | 1.3 ± 0.3 | 1.8 ± 0.3* | 2.0 ± 0.4** | 1.3 ± 0.3 |
| DAP | 69.6 ± 12.1 | 68.3 ± 17.9 | 64.7 ± 8.0 | 65.9 ± 14.7 |
| Spermine | 16.1 ± 0.8 | 16.2 ± 1.9 | 21.2 ± 1.2** | 15.5 ± 1.8 |
| Spermidine | 112.3 ± 7.3 | 154.5 ± 37.9* | 144.8 ± 21.9* | 115.0 ± 25.0 |
| Putrescine | 275.8 ± 58.1 | 307.2 ± 34.5 | 189.4 ± 15.4* | 178.1 ± 37.5* |
| Ornithine | 146.3 ± 17.0 | 136.5 ± 21.9 | 127.9 ± 20.1 | 133.8 ± 21.8 |
| Arginine | 188.2 ± 24.5 | 189.1 ± 46.1 | 143.8 ± 31.7* | 183.2 ± 43.2 |
