## Supplementary Fig S1 for "Insight into GABA shunt-associated aldehyde dehydrogenases (ALDH) and stress responses of ALDH superfamily in moss and barley"

**Supplementary Fig. S1. Gene annotation corrections and relative expression levels of *ALDH* superfamily genes in *Physcomitrium patens* and *Hordeum vulgare*. (A)** Exon 9 of *PpALDH11A5*. An annotated gene Pp6c17_180 lacks a part of exon 9, while the whole exon is indeed present in the moss genome and comprises a stop codon. **(B)** Three copies of *HvALDH2B5* on chromosome 6 in *H. vulgare*. Two gene annotations covering the region are in red. **(C)** Relative expression levels in moss normalized to *PpALDH5F2*. **(D)** Relative expression levels in barley normalized to *HvALDH5F1*. Data represent the means ± S.D. (n = 6).

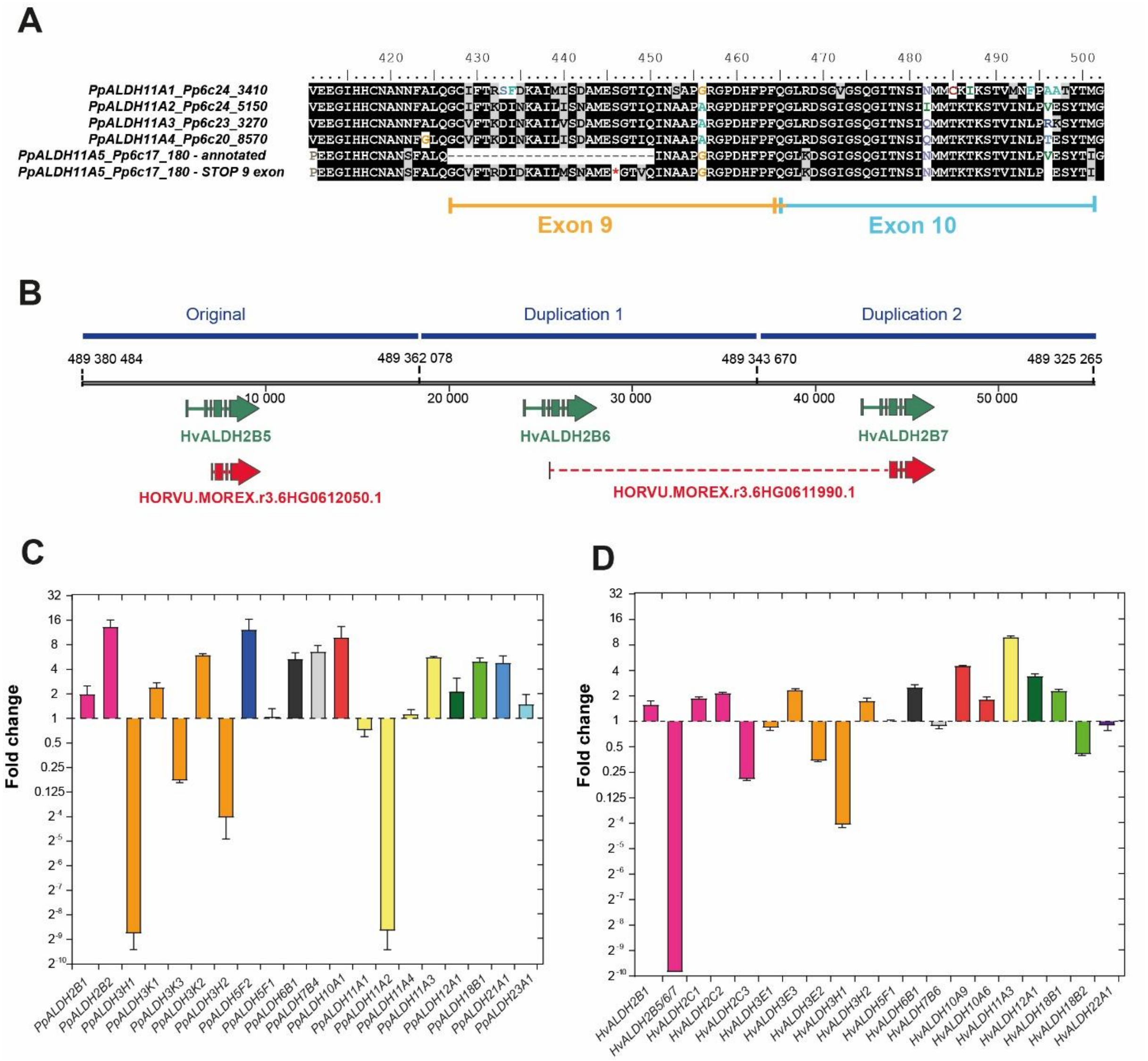
