## Supplementary Fig S2 for "Insight into GABA shunt-associated aldehyde dehydrogenases (ALDH) and stress responses of ALDH superfamily in moss and barley"

**Supplementary Fig. S2. Promoter analysis of *cis*-regulatory elements.** The table shows several identified sites associated with hormone responses in blue, such as ABA-Responsive Element (ABRE), auxin-responsive region (AuxRR) core, CGTCA-motif responsive to jasmonate and ABA, gibberellin-responsive element P-box, salicylate-responsive elements TCA-element and as-1, MeJA-responsive TGACG-motif/TGTCA-element, GARE-motif (Gibberellin-Activated REsponse), and Ethylene-Responsive Element (ERE). Elements associated with general stress are in green and include Wound-Responsive Elements (WRE) and WUN-motif, Anaerobic Response Element (ARE), MYB-binding site (MBS), drought-responsive MYB site, MYC-site, Stress-Responsive Element (STRE), pathogen-related W-box, Low-Temperature Responsive element (LTR), TC-rich repeats and dehydration-responsive element (DRE).

| Gene | ABRE | AuxRR-core | CGTCA motif | P-box | TCA element | TGACG motif | as-1 | GARE motif | TGA element | ERE | WRE | ARE | MBS | MYB site | MYC site (bHLHZip) | STRE | W box | LTR | TC-rich repeats | DRE motif | WUN motif | Hormone response | Stress Response |
|---|---|---|---|---|---|---|---|---|---|---|---|---|---|---|---|---|---|---|---|---|---|---|---|
| *PpALDH2B1* | 0 | 0 | 0 | 0 | 0 | 0 | 0 | 0 | 0 | 0 | 1 | 2 | 0 | 3 | 0 | 0 | 1 | 0 | 0 | 0 | 0 | 0 | 7 |
| *PpALDH2B2* | 5 | 0 | 5 | 0 | 0 | 5 | 5 | 0 | 1 | 0 | 0 | 0 | 3 | 5 | 1 | 0 | 0 | 0 | 1 | 0 | 0 | 21 | 10 |
| *PpALDH3H1* | 2 | 0 | 4 | 0 | 1 | 4 | 4 | 1 | 0 | 0 | 1 | 1 | 0 | 3 | 1 | 3 | 0 | 0 | 0 | 0 | 0 | 16 | 9 |
| *PpALDH3K1* | 1 | 0 | 1 | 1 | 0 | 1 | 1 | 0 | 0 | 0 | 2 | 1 | 0 | 3 | 0 | 0 | 0 | 1 | 0 | 0 | 0 | 5 | 7 |
| *PpALDH3K3* | 2 | 0 | 1 | 0 | 0 | 1 | 1 | 1 | 0 | 0 | 0 | 1 | 2 | 8 | 0 | 0 | 0 | 0 | 0 | 0 | 0 | 6 | 11 |
| *PpALDH3K2* | 1 | 1 | 1 | 1 | 0 | 1 | 1 | 0 | 0 | 0 | 0 | 0 | 0 | 1 | 3 | 1 | 0 | 0 | 0 | 0 | 0 | 6 | 5 |
| *PpALDH3H2* | 1 | 0 | 1 | 0 | 0 | 1 | 1 | 0 | 2 | 0 | 0 | 3 | 0 | 3 | 1 | 1 | 1 | 0 | 0 | 0 | 1 | 6 | 10 |
| *PpALDH5F2* | 1 | 0 | 2 | 0 | 2 | 2 | 2 | 0 | 0 | 0 | 2 | 1 | 0 | 0 | 2 | 2 | 0 | 0 | 1 | 0 | 1 | 9 | 9 |
| *PpALDH5F1* | 4 | 0 | 2 | 0 | 1 | 1 | 1 | 0 | 0 | 0 | 2 | 1 | 0 | 4 | 3 | 1 | 0 | 0 | 0 | 1 | 0 | 9 | 12 |
| *PpALDH6B1* | 2 | 0 | 5 | 1 | 2 | 5 | 5 | 1 | 0 | 0 | 1 | 1 | 0 | 2 | 1 | 1 | 0 | 0 | 0 | 1 | 0 | 21 | 7 |
| *PpALDH7B4* | 2 | 0 | 3 | 0 | 1 | 3 | 3 | 0 | 1 | 0 | 3 | 0 | 2 | 6 | 2 | 5 | 0 | 0 | 1 | 0 | 0 | 13 | 19 |
| *PpALDH10A1* | 1 | 0 | 1 | 0 | 1 | 1 | 1 | 0 | 2 | 2 | 0 | 1 | 0 | 1 | 4 | 1 | 0 | 1 | 0 | 0 | 0 | 9 | 8 |
| *PpALDH11A1* | 0 | 0 | 3 | 0 | 0 | 3 | 3 | 1 | 1 | 0 | 1 | 1 | 2 | 6 | 0 | 2 | 2 | 1 | 0 | 2 | 0 | 11 | 17 |
| *PpALDH11A2* | 3 | 0 | 3 | 1 | 1 | 3 | 3 | 0 | 1 | 0 | 1 | 3 | 0 | 2 | 1 | 2 | 0 | 1 | 0 | 1 | 0 | 15 | 11 |
| *PpALDH11A4* | 0 | 0 | 2 | 0 | 1 | 2 | 2 | 0 | 2 | 0 | 1 | 2 | 2 | 6 | 0 | 1 | 0 | 0 | 0 | 0 | 0 | 9 | 12 |
| *PpALDH11A3* | 1 | 0 | 2 | 0 | 0 | 2 | 2 | 1 | 0 | 0 | 0 | 2 | 1 | 5 | 2 | 1 | 0 | 0 | 0 | 0 | 0 | 8 | 11 |
| *PpALDH12A1* | 2 | 0 | 3 | 0 | 0 | 3 | 3 | 1 | 1 | 0 | 0 | 2 | 1 | 4 | 3 | 3 | 2 | 0 | 0 | 0 | 0 | 13 | 15 |
| *PpALDH18B1* | 0 | 0 | 1 | 0 | 0 | 1 | 1 | 0 | 0 | 0 | 0 | 0 | 0 | 2 | 1 | 0 | 0 | 0 | 0 | 1 | 0 | 3 | 4 |
| *PpALDH21A1* | 0 | 0 | 3 | 1 | 2 | 3 | 3 | 1 | 1 | 2 | 0 | 1 | 0 | 1 | 4 | 1 | 0 | 0 | 0 | 0 | 0 | 16 | 7 |
| *PpALDH23A1* | 0 | 0 | 0 | 0 | 1 | 0 | 0 | 0 | 0 | 0 | 0 | 2 | 1 | 4 | 1 | 0 | 0 | 0 | 0 | 0 | 0 | 1 | 8 |
| *HvALDH2B1* | 6 | 0 | 1 | 0 | 0 | 1 | 1 | 1 | 0 | 0 | 0 | 1 | 0 | 5 | 0 | 2 | 0 | 0 | 0 | 0 | 0 | 10 | 8 |
| *HvALDH2B5* | 1 | 0 | 2 | 0 | 0 | 1 | 2 | 1 | 0 | 0 | 0 | 1 | 0 | 2 | 2 | 0 | 0 | 0 | 0 | 1 | 0 | 7 | 6 |
| *HvALDH2B6* | 1 | 0 | 2 | 0 | 0 | 1 | 2 | 1 | 0 | 0 | 0 | 1 | 0 | 2 | 2 | 0 | 0 | 0 | 0 | 1 | 0 | 7 | 6 |
| *HvALDH2B7* | 1 | 0 | 2 | 0 | 0 | 1 | 2 | 1 | 0 | 0 | 0 | 1 | 0 | 2 | 2 | 0 | 0 | 0 | 0 | 1 | 0 | 7 | 6 |
| *HvALDH2C1* | 5 | 0 | 0 | 0 | 1 | 0 | 0 | 0 | 1 | 0 | 1 | 1 | 0 | 1 | 2 | 4 | 1 | 0 | 0 | 0 | 0 | 7 | 9 |
| *HvALDH2C2* | 6 | 0 | 4 | 0 | 0 | 4 | 4 | 0 | 1 | 0 | 1 | 1 | 1 | 10 | 1 | 1 | 1 | 0 | 2 | 2 | 0 | 19 | 19 |
| *HvALDH2C3* | 3 | 0 | 1 | 0 | 0 | 2 | 2 | 0 | 1 | 0 | 1 | 1 | 1 | 5 | 0 | 2 | 0 | 0 | 0 | 4 | 0 | 9 | 13 |
| *HvALDH3E1* | 4 | 0 | 1 | 0 | 0 | 1 | 1 | 0 | 0 | 0 | 0 | 0 | 0 | 4 | 2 | 1 | 0 | 0 | 0 | 0 | 0 | 7 | 7 |
| *HvALDH3E3* | 3 | 0 | 0 | 1 | 0 | 0 | 0 | 0 | 0 | 0 | 0 | 2 | 1 | 3 | 1 | 5 | 0 | 2 | 0 | 0 | 0 | 4 | 14 |
| *HvALDH3E2* | 3 | 0 | 3 | 0 | 1 | 3 | 3 | 0 | 0 | 1 | 0 | 0 | 0 | 3 | 2 | 0 | 0 | 0 | 2 | 0 | 1 | 14 | 8 |
| *HvALDH3H1* | 5 | 0 | 0 | 1 | 0 | 0 | 0 | 0 | 0 | 0 | 2 | 1 | 0 | 2 | 0 | 1 | 0 | 1 | 0 | 1 | 0 | 6 | 6 |
| *HvALDH3H2* | 4 | 1 | 2 | 0 | 1 | 2 | 2 | 0 | 0 | 0 | 0 | 2 | 0 | 3 | 2 | 2 | 0 | 0 | 0 | 0 | 0 | 12 | 9 |
| *HvALDH5F1* | 4 | 0 | 1 | 0 | 0 | 1 | 1 | 0 | 1 | 0 | 0 | 1 | 0 | 7 | 4 | 3 | 0 | 0 | 0 | 1 | 0 | 8 | 16 |
| *HvALDH6B1* | 3 | 0 | 5 | 0 | 0 | 4 | 4 | 0 | 0 | 2 | 1 | 0 | 0 | 2 | 1 | 3 | 0 | 0 | 0 | 0 | 0 | 18 | 6 |
| *HvALDH7B6* | 5 | 0 | 4 | 0 | 0 | 5 | 5 | 0 | 0 | 0 | 4 | 0 | 0 | 2 | 1 | 4 | 1 | 0 | 0 | 2 | 0 | 19 | 10 |
| *HvALDH10A9* | 4 | 0 | 4 | 0 | 0 | 5 | 5 | 2 | 0 | 0 | 0 | 2 | 0 | 6 | 1 | 3 | 1 | 0 | 0 | 0 | 0 | 20 | 13 |
| *HvALDH10A6* | 5 | 0 | 4 | 0 | 0 | 4 | 4 | 0 | 1 | 2 | 2 | 0 | 0 | 0 | 0 | 2 | 0 | 0 | 0 | 1 | 0 | 20 | 3 |
| *HvALDH11A3* | 6 | 1 | 2 | 0 | 0 | 2 | 2 | 0 | 0 | 0 | 0 | 0 | 1 | 6 | 0 | 1 | 0 | 0 | 0 | 1 | 0 | 13 | 9 |
| *HvALDH12A1* | 1 | 0 | 2 | 0 | 0 | 1 | 2 | 0 | 0 | 0 | 0 | 1 | 0 | 1 | 1 | 1 | 0 | 0 | 1 | 1 | 1 | 6 | 7 |
| *HvALDH18B1* | 5 | 0 | 4 | 0 | 0 | 3 | 4 | 0 | 0 | 0 | 1 | 0 | 0 | 1 | 1 | 4 | 0 | 1 | 1 | 1 | 0 | 16 | 9 |
| *HvALDH18B2* | 4 | 0 | 0 | 0 | 0 | 0 | 0 | 0 | 0 | 0 | 3 | 1 | 0 | 1 | 1 | 2 | 2 | 0 | 0 | 0 | 0 | 4 | 7 |
| *HvALDH22A1* | 9 | 0 | 0 | 0 | 0 | 0 | 0 | 0 | 0 | 1 | 0 | 0 | 0 | 2 | 0 | 4 | 0 | 1 | 0 | 2 | 0 | 10 | 9 |
