## Supplementary Fig S3 for "Insight into GABA shunt-associated aldehyde dehydrogenases (ALDH) and stress responses of ALDH superfamily in moss and barley"

**Supplementary Fig. S3. Gel permeation chromatography of moss ΔPpALDH10A1 and PpALDH5F1. (A)** ΔPpALDH10A1 migrates as a dimer on the Superdex 200 (10/300) column. Gel permeation chromatography was performed in 50 mM phosphate buffer, pH 7.0, with 150 mM NaCl. **(B)** PpALDH5F1 migrates as a tetramer on the Superdex 200 (10/300) column.

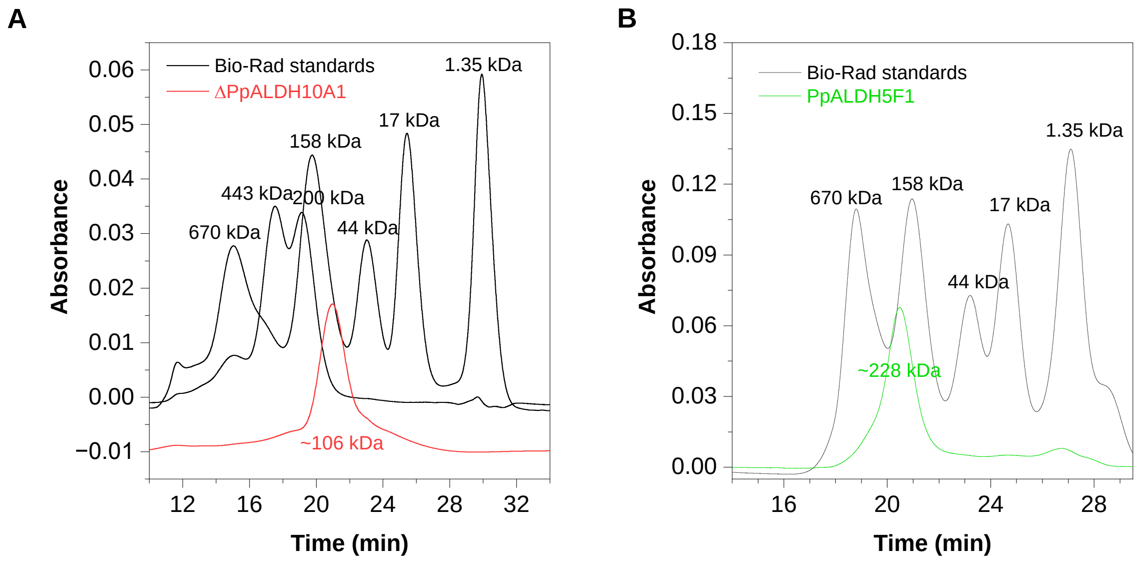
