## Supplementary Fig S4 for "Insight into GABA shunt-associated aldehyde dehydrogenases (ALDH) and stress responses of ALDH superfamily in moss and barley"

**Supplementary Fig. S4. Sequence alignment of three plant ALDH5 enzymes analyzed in this work with human ALDH5A1.** The active site residues studied by site-directed mutagenesis are labeled. Sequences used: human ALDH5A1 (Uniprot P51649.2), PpALDH5F1 (A0A2K1ICJ7), PpALDH5F2 (A9TPC4) and HvALDH5F1 (M0XSP9).

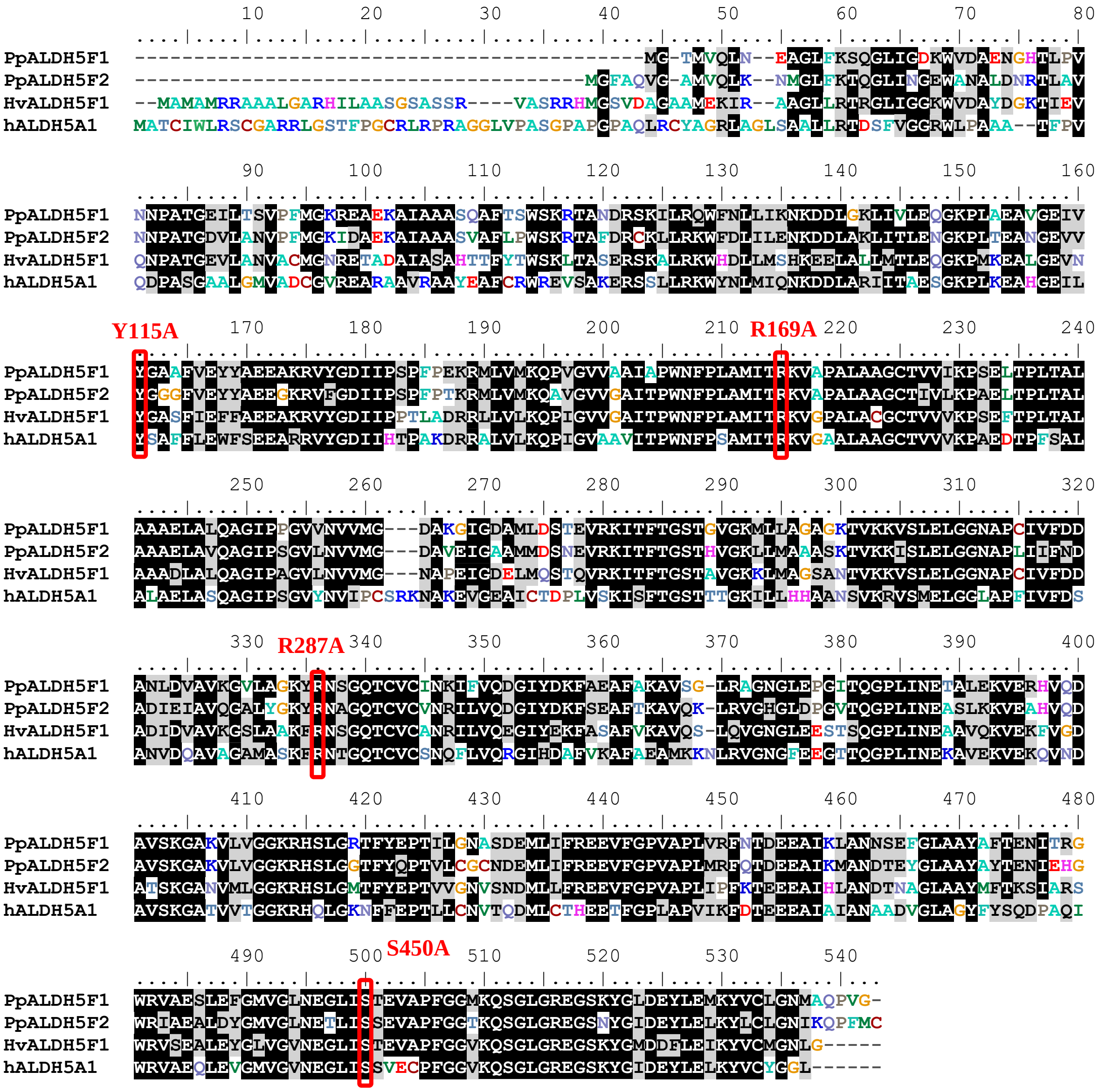
