## Supplementary Fig S5 for "Insight into GABA shunt-associated aldehyde dehydrogenases (ALDH) and stress responses of ALDH superfamily in moss and barley"

**Supplementary Fig. S5. Example of a construct preparation and genotyping of *Physcomitrium aldh* knockouts. (A)** Construct preparation for homologous recombination. Cloned 1000 bp fragments are colored in yellow (labelled as cassette 1 and 2), the resistance marker (colored in pink) is placed behind the 35S promoter between both cassettes. Cloning and sequencing primers are labelled red. **(B)** PCR screening of *Physcomitrium* knockout mutants for the presence of the complete cassette, including fragments 1 and 2, and confirmation of the correct recombination event. The red arrow indicates the expected size. **(C)** Confirmation of the absence of *ALDH* transcripts in *Physcomitrium* knockout mutants (*versus* WT) by qPCR in 35 cycles using specific FAM-TAM probes.

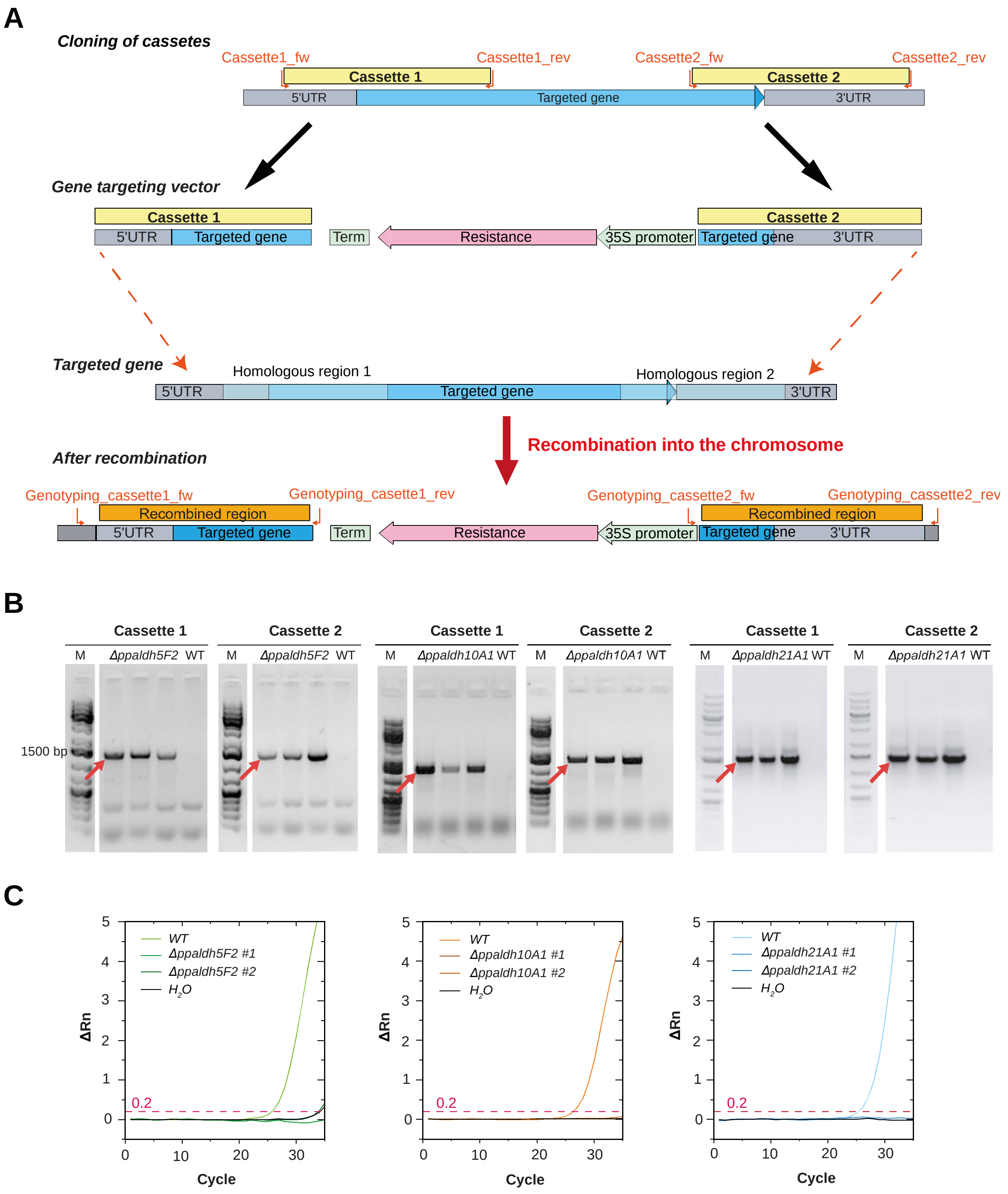
