## Supplementary Fig S6 for "Insight into GABA shunt-associated aldehyde dehydrogenases (ALDH) and stress responses of ALDH superfamily in moss and barley"

**Supplementary Fig. S6. Volcano plots illustrating differentially expressed genes in three moss *aldh* mutants vs WT in RNA-seq data.** Each point represents a gene, with the x-axis displaying the $\log_2$ fold change and the y-axis showing statistical significance as $-\log_{10}$ of q-value. Significantly upregulated and downregulated genes ($q \leq 0.05$, fold-change $\geq 2$) are highlighted in green for *aldh5F2*, blue for *aldh21A1* and orange for *aldh10A1*, while non-significant genes are shown in black.

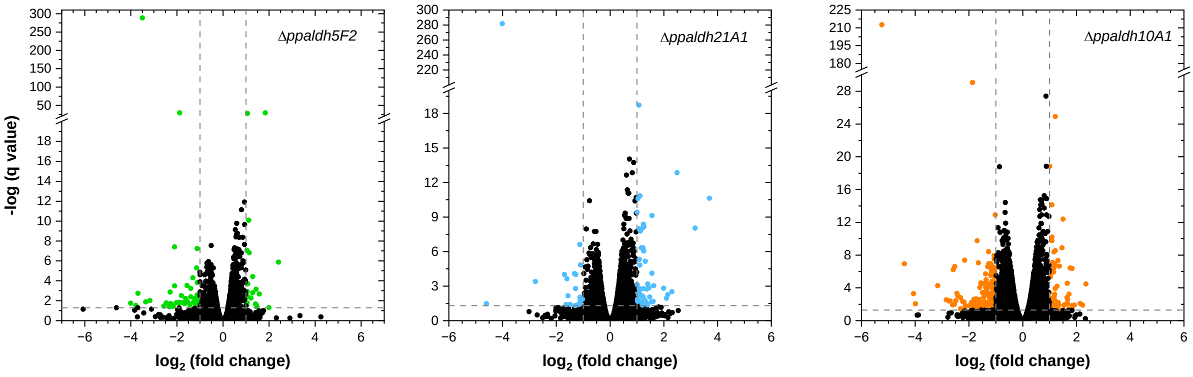
