## Supplementary Fig S7 for "Insight into GABA shunt-associated aldehyde dehydrogenases (ALDH) and stress responses of ALDH superfamily in moss and barley"

**Supplementary Fig. S7. A heat map of DEGs in moss *aldh5F2* mutant vs control generated from RNA seq data reflecting log(e)-transformed transcripts per million (TPM) values.**

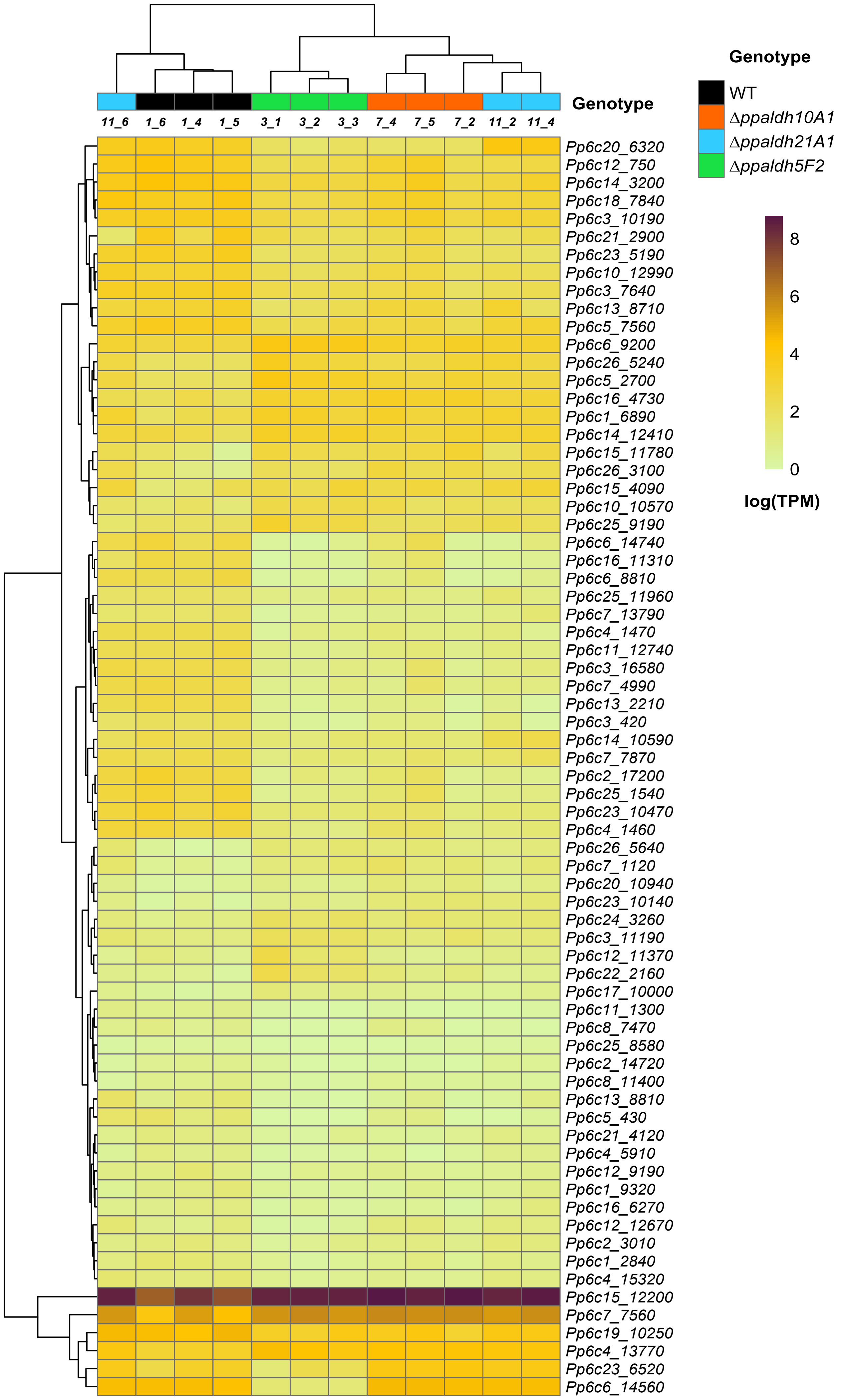
