## Supplementary Fig S8 for "Insight into GABA shunt-associated aldehyde dehydrogenases (ALDH) and stress responses of ALDH superfamily in moss and barley"

**Supplementary Fig. S8. A heat map of DEGs in moss *aldh21A1* vs control generated from RNA seq data reflecting log(e)-transformed transcripts per million (TPM) values.**

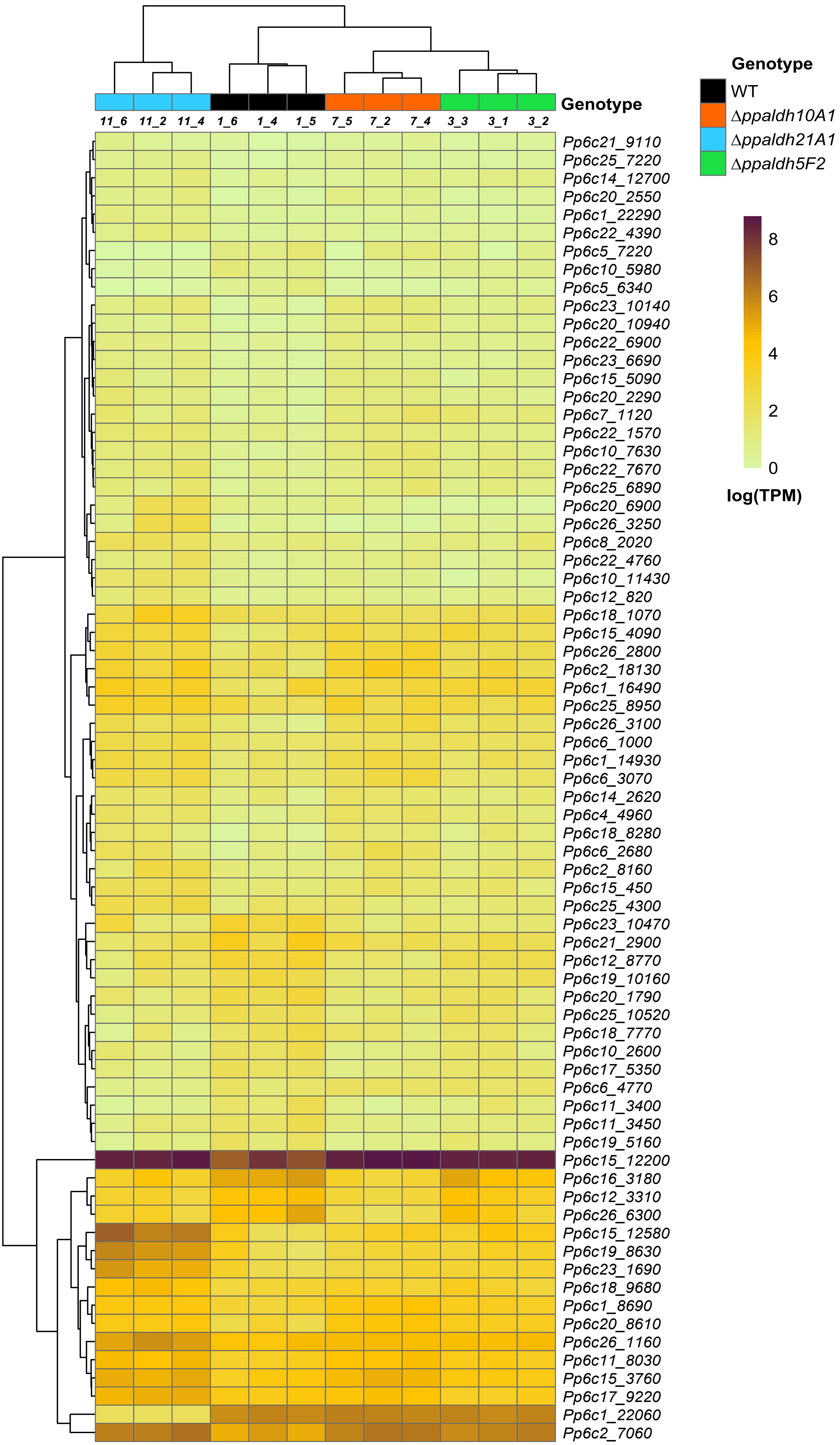
