## Supplementary Fig S10 for "Insight into GABA shunt-associated aldehyde dehydrogenases (ALDH) and stress responses of ALDH superfamily in moss and barley"

**Supplementary Fig. S10. Photosynthetic control parameters in moss *aldh* mutants during light exposure. (A)** The effective quantum yield of photosystem I (PSI) photochemistry Y(I). **(B, C)** Quantum yields of non-photochemical energy dissipation in PSI due to donor (Y(ND)) and acceptor side limitation (Y(NA)), respectively. **(D)** Non-photochemical quenching (NPQ) in PSII. Data were recorded in cultures dark-adapted for 30 min and then exposed to actinic light (278 µmol photons $m^{-2}$ $s^{-1}$). The parameters were calculated using saturating red-light pulses (300 ms, 10,000 µmol photons $m^{-2}$ $s^{-1}$) during actinic light exposure, followed by a dark relaxation phase. **(E)** $F_V/F_M$ ratio representing a maximal quantum yield of the photosystem II (PSII) photochemistry. Values are means ± S.D. (n=7)

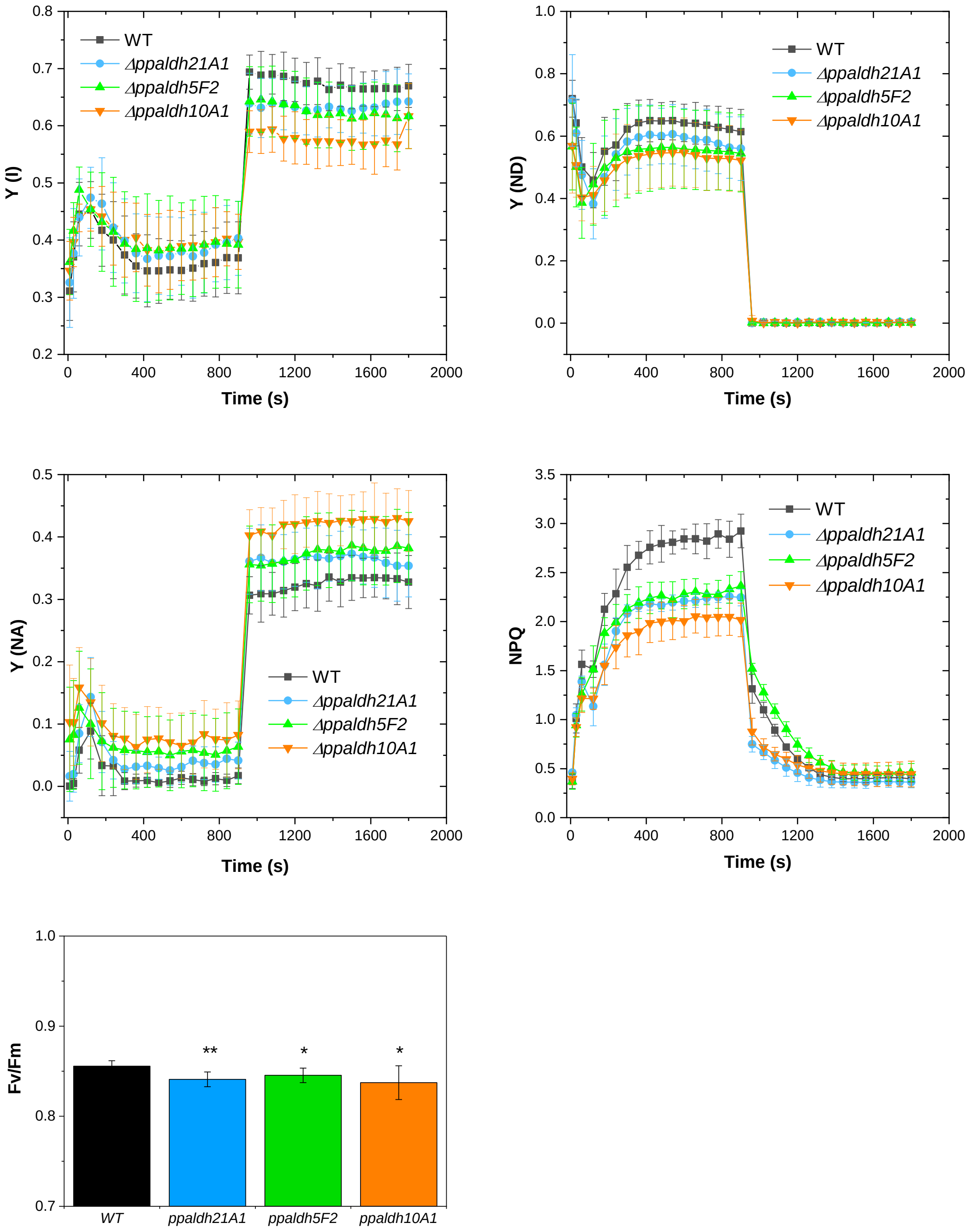
